# Proteomics for cultivated meat: the importance of Analytical Standardization

**DOI:** 10.64898/2026.03.23.713501

**Authors:** João Palma, Chloe Colchero Leblanc, Remy Kusters, Franks Kamgang Nzekoue

## Abstract

Cultivated meat production requires robust and validated analytical methods for comprehensive characterization. While transcriptomics-based approaches establish the foundational profile of molecular analysis, proteomics provides additional resolution that further enhances scientific certainty in both product development and safety characterization. However, the industry adoption of proteomics is currently hindered by technical complexity and a critical lack of analytical standardization, which leads to significant workflow-dependent variations in proteome coverage. To address this gap, we investigated the influence of key workflow steps (digestion, cleanup, LC-MS conditions) on the proteome profile of cultivated duck biomass. We compared five bottom-up sample preparation protocols – two traditional in-solution options (urea and SDC-based protocols), two device-based approaches (PreOmics iST and EasyPep kits), and an innovative protocol (SPEED), and demonstrated that device-based protocols offered the highest peptide yield and proteome coverage. However, optimization allowed cost-effective in-solution methods to achieve comparable performance. Specifically, an optimal digestion time of 3 hours at 37°C and the use of polymer-based desalting columns significantly enhanced protein identification (∼4500 - 5000 IDs). Moreover, data independent acquisition (DIA) provided deeper proteome coverage than data dependent acquisition (DDA) with higher precision (∼6500 vs 5000 IDs). The validated Standard Operating Procedures presented here establish a standardized framework for bulk bottom-up proteomics in cultivated meat, facilitating the generation of reliable and comparable data required for robust multi-omics characterization.

**Graphical abstract:** 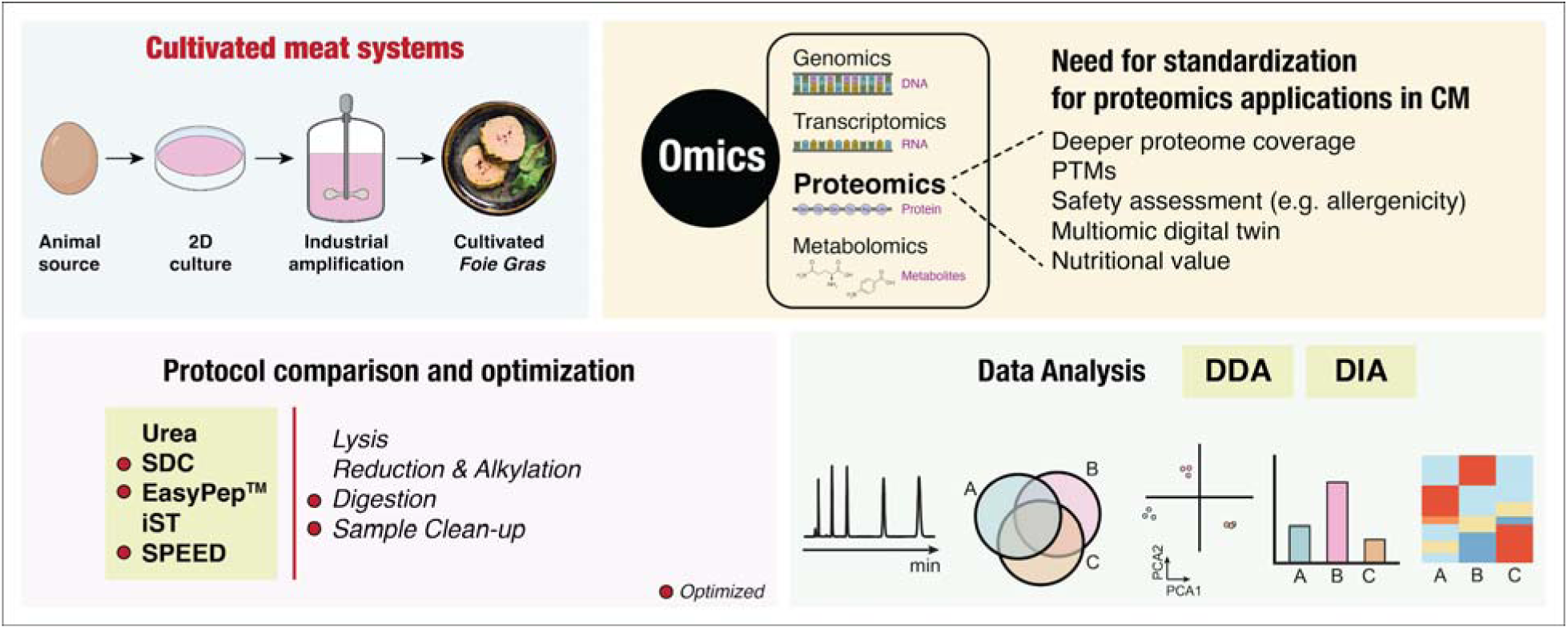

**Highlights:**

1. Complexity and non-standardization limit MS-proteomics use in cultivated meat (CM).
2. CM protein profile varies with sample prep, LC-MS, and data processing pipeline.
3. Device-based and optimized cost-effective protocols offer a high proteome coverage.
4. Proteomics can complement transcriptomics for a comprehensive CM characterization.
5. Proposed standardized methods ensure reliable data for future regulatory submissions.

## Introduction

Cultivated meat is a sustainable and promising alternative to traditionally produced meat, driven by advancements in bioprocessing, media cost reduction, and industrial scaling. This innovation contributes to the global protein transition aiming to tackle the current food system challenges, including global demographic expansion, animal welfare concerns, and environmental exigencies (Kardas et al. 2025; Kim et al., 2025; Singh et al., 2026).

With the emergence of cultivated meat, proteomics - the large-scale study of proteins, their structure and function - can be considered as a valuable analytical asset to further build consumer confidence (Lee & Hur, 2025; Gu et al., 2025). Indeed, given the core mission of cultivated meat production to provide an alternative protein source, a thorough characterization of its proteome across the entire cultivation workflow holds significant analytical value. Proteomics can serve to rigorously evaluate the effect of scaling and processes on protein quality, demonstrate genetic stability, allergenicity, cell-line authenticity, and ensure nutritional and organoleptic parity with conventional meat.

Moreover, proteomics is a powerful tool for understanding cellular biology (Eldrid et al., 2025). Integrating proteomics into a multi-omics strategy can further enhance the comprehensive characterization of cultivated meat across various biological scales (Mathieu et al., 2025). This multi-omics approach is relevant in key research areas such as cell differentiation, media development (Trautmann et al., 2025), and for digital twin modeling allowing *in-silico* control and optimization of cultured meat production (Qiu et al., 2026). Furthermore, it is particularly valuable within New Approach Methodologies (NAMs), which regulatory agencies increasingly encourage for the safety assessment of novel foods like cultivated meat (Levine et al., 2025; Deng et al. 2025). Specifically, transcriptomics are currently used to demonstrate e.g. the absence of mRNA expression for allergenic proteins or the absence of upregulation of known allergens in the final cultured product. Integrating a proteomic layer into a comprehensive multi-omics digital twin, can strengthen current NAMs to provide an alternative to animal studies in safety assessment (Prabahar et al., 2024). This shift toward NAMs and replacing *in vivo* animal testing aligns with the principles of the 3Rs (reduce, refine, and replace animal studies) and the ultimate goal of the European Union to avoid the use of live animals for scientific testing as soon as it is scientifically possible (Directive 2010/63/EU).

Despite its potential and the technical advancements linked to Mass Spectrometry (MS), the application of proteomics in the industry still faces significant challenges due to its inherent complexity and the lack of standardization. Indeed, according to the goals of analysis - targeted or untargeted proteomics, Post-translational modifications (PTMs)/proteoforms or canonical proteome, single-cell or spatial proteomics - the workflow required is highly specific and can be easily disrupted (Guo et al., 2025; Hu & Wang, 2024). Moreover, unlike genomics and transcriptomics, universal proteomics protocols with high robustness are currently unavailable, especially in the field of cultivated meat where it can play a major role (Ham et al., 2025; Ledwith et al., 2024). In addition, while modern transcriptomics like next-generation sequencing (RNA seq) offer a comprehensive coverage by sequencing all expressed genes, high-throughput proteomics provide a more limited view of the proteome, which is constrained by the specific analytical workflow employed (Karnaneedi et al., 2023; Molho et al., 2024). Indeed, for an analytical strategy such as label-free quantitative proteomics, there is an overwhelming number of protocols in the literature with variations in sample preparation, data acquisition strategies (DDA, DIA) and processing workflows. These variations represent a significant gap in our current knowledge regarding how they could ultimately influence the proteome coverage of cultivated meat (Woodland et al., 2025).

To overcome these challenges and ensure reliable and comparable data, the standardization of proteomic methods is essential. Bottom-up proteomics for bulk cell analysis is the most widely adopted approach for proteome analysis to date. Therefore, this workflow should be prioritized for standardization in cultivated meat characterization. In this approach, proteins are enzymatically digested into peptides, which are then analyzed by LC-MS (Varnavides et al., 2022). The resulting peptide MS spectra are matched against protein databases, allowing the original proteins to be identified and quantified. Because peptides are easier to separate, fragment, and identify compared to whole proteins (top-down approach), bottom-up proteomics enables high-throughput identification and quantification of thousands of proteins from complex biological samples (Kaulich & Tholey, 2025). The results of this approach ultimately depend on the quality of three major stages: (1) sample preparation, (2) LC-MS acquisition, and (3) computational data processing and analysis (Kanao et al., 2024). However, sample preparation alone involves multiple critical steps, including protein extraction, reduction, alkylation, enzymatic digestion, peptide cleanup and enrichment, all of which can significantly influence the quality and depth of the resulting proteome (Schar et al., 2025).

To our knowledge, this study is the first to address the critical need for standardized proteomics approaches in cultivated meat, aiming to establish new standards and bring proteomics to the next level for industry development and global regulatory adoption. In order to reduce the knowledge gap regarding the influence of workflow variability on the cultivated meat proteome, we systematically compared the most referenced bottom-up sample preparation protocols. Moreover, by investigating the impact of key conditions - detergent, protein digestion, cleanup strategy, LC-MS conditions - on the proteome coverage, this study ultimately presents optimized bottom-up protocols specifically tailored for cultivated meat application.

## 2. Materials and methods

### 2.1. Cell culture and sample collection

Cultivated meat biomass from duck cell culture (*A. platyrhynchos* (Pekin duck) cells) was produced by PARIMA in suspension bioreactors under sterile conditions following its proprietary serum-free media, bioprocess, and downstream technology. For the purpose of comparing different protocols, several aliquots from the same production batch were prepared and stored (−80°C), each containing a consistent amount of protein.

### 2.2. Experiment design and comparison of sample preparation protocols

The first study aimed to review, select, and compare the main referenced and established protocols for sample preparation in label-free bottom-up proteomics. Five different protocols were thus selected based on their frequency of use in the literature and potential for standardization:

– Two traditional protocols (in solution-digestion): with nonionic chaotropic agent (urea - Method 1) and ionic detergent (Sodium deoxycholate - Method 2).
– Two device-based protocols: EasyPep™ and PreOmics iST, which offer streamlined protocols with pre-optimized reagents (Methods 3 and 4 respectively).
– One innovative in-solution protocol (SPEED - Sample Preparation by Easy Extraction and Digestion, Method 5).

#### 2.2.1. Method 1. In-Solution protocol – Urea-Based extraction

Cultivated biomass samples stored at −80°C were thawed and lysed with 8 M urea (in 0.1 M Tris-HCl, pH 8.5). After centrifugation, protein concentration was determined using a Pierce BCA Protein Assay Kit (ThermoFisher Scientific). Proteins were then reduced with 10 mM Dithiothreitol (DTT) for 1 h at 37 °C, followed by alkylation with 20 mM Iodoacetamide (IAA) for 30 min in the dark at room temperature (RT). The sample was diluted 1:8 to reduce the urea concentration to 1 M using 50mM Tris-HCl (pH 8). Finally, trypsin/LysC was added at a 1:20 enzyme-to-protein ratio and incubated overnight at 37 °C. Digestion was stopped by adding Trifluoroacetic acid (TFA) to a final concentration of 1%, and the digested peptide was desalted using C18 tips, dried in a SpeedVac concentrator, and resuspended in 0.1% formic acid (FA) for LC-MS analysis.

#### 2.2.2. Method 2. In-Solution Protocol – SDC-Based extraction

Biomass was lysed with 1% Sodium deoxycholate (SDC, Sigma-Aldrich) in 0.1 M Tris-HCl (pH 8.5). The lysate was clarified by centrifugation, reduced with 10 mM DTT for 1 h at 60 °C, and then alkylated with 20 mM IAA for 30 min in the dark at RT. Proteins were digested overnight at 37 °C with trypsin/LysC at a 1:20 ratio. Digestion was stopped with TFA (to a final concentration of 1%) and SDC precipitates were removed by centrifugation (14.000 g, 1 min). Peptides were desalted using C18 tips, dried, and resuspended in 0.1% FA for LC-MS analysis.

#### 2.2.3. Method 3. Device-based protocol – EasyPep™

Sample preparation was performed following the manufacturer instructions (Thermo Scientific™ EasyPep™ 96 MS Sample Prep Kit User Guide). Briefly, after biomass lysis, extracted protein was simultaneously reduced and alkylated at 95 °C for 10 min using the proprietary reagent mix. Digestion was carried out with Trypsin/Lys-C mix for 3 h at 37 °C. Peptide desalting was performed using the peptide cleanup columns included in the kit. After elution and drying, peptides were resuspended in 0.1% FA for LC-MS analysis.

#### 2.2.4. Method 4. Device-based protocol – In-StageTip sample prep

Biomass was lysed and digested using the iST sample kit according to manufacturer protocol (PreOmics). Briefly, biomass was transferred into a StageTip and processed in three steps: (1) lysis, denaturation, reduction, and alkylation occurring in an appropriate buffer for 10 min at 95°C; (2) digestion with Trypsin/LysC at 37 °C for 3 h; (3) peptide purification using an appropriate cartridge. The resulting peptides are dried and resuspended in 0.1% FA.

#### 2.2.5. Method 5. SPEED (Sample Preparation by Easy Extraction and Digestion)

Biomass was initially resuspended in trifluoroacetic acid (TFA) at a 1:4 (sample-to-TFA, v/v) ratio and left to incubate for 5 minutes at RT. Subsequent neutralization of the acid was achieved by adding 2 M Tris base (8x the volume of TFA originally used). Protein was reduced and alkylated with 10 mM Tris(2-carboxyethyl)phosphine (TCEP) and 40 mM Chloroacetamide (CAA) for 5 minutes at 95 °C. The sample was diluted with MilliQ water (1:5), and trypsin was added at a 1:20 enzyme-to-protein ratio. Digestion was carried out overnight at 37 °C. After digestion, the reaction was stopped by adding TFA to a 2% final concentration, and peptides were desalted using C18 tips, dried, and resuspended for LC-MS.

### 2.3. Effect of digestion conditions

Following the selection of most promising protocols, a study was undertaken to evaluate how digestion conditions affect proteomics results. This investigation considered the variability in protocols regarding the digestion time (from 1 h to overnight) and the emerging protocols proposing higher temperatures (e.g., 47°C) to significantly reduce the overall sample preparation time. Five digestion conditions were thus compared: 1 h at 47 °C, 1h at 37°C, 2h at 37°C, 3h at 37°C, and 18 h at 37 °C (for overnight conditions).

### 2.4. Impact of peptide cleanup strategy

To evaluate the impact of different cleanup strategies on peptide recovery and proteome coverage, three cleanup methods were tested using different cleanup columns: C18 StageTips (C18 reversed-phase tips), Pierce™ C18 Spin Columns (C18 reversed-phase column), and Pierce™ Peptide Desalting Spin Columns (with polymer-based hydrophobic resin). The Peptide yield and the proteome coverage was compared across the cleanup approaches.

### 2.5. Impact of analytical workflow

The impact of the LC-MS acquisition workflow was also studied comparing proteome profiles obtained through Data-Dependant acquisition (DDA) and Data-Independant acquisition (DIA). Indeed, if DDA is the reference method for labeled and label-free quantification, there is a significant interest towards DIA in the proteomics community. Moreover, the impact of the peptide loading amount (0.5 - 6 ug) was also assessed.

### 2.6. Liquid chromatography-mass spectrometry (LC-MS)

LC-MS analyses were performed using a Vanquish Neo™ UHPLC system coupled to an Orbitrap Exploris™ 480 MS (Thermo Fisher Scientific). Peptides were separated in a trap and elute mode using a PepMap Neo C18 (75 μm × 500 mm, 2 μm C18; Thermo Fisher Scientific) at a temperature of 50°C. The mobile phase consisted of ultrapure water (A) and 80 % ACN (B) both containing 0.1 % FA at a flow rate of 300 nL/min. The LC gradient was 0 - 35% B over 60 min, 35 - 95% B in 10 s, and 95% for 4 min. The flow rate ramped from 300 to 450 nL/min over 64 min. Peptide analysis was performed in POS through DDA and DIA modes. DDA was performed through the following parameters: MS1 full scan in each scan cycle (R= 120 K, AGC= 300 %, max IT= 50 ms, RF Lens (%)= 40, scan range= 350–1400 m/z) followed by ddMS2 scans (R= 15 K, AGC= Standard, max IT= auto, cycle time = 2 s). The higher-energy collisional dissociation (HCD) was set to 30 %, and the precursor isolation width was 1.6 m/z. A dynamic exclusion window of 45 s was used in this study. DIA parameters were as follow: MS1 (R= 120 K, AGC= 300 %, max IT= 50 ms, RF Lens (%)= 40, scan range= 350–1400 m/z) with DIAScan (R= 30 K, Isolation window= 15 m/z, Window Overlap (m/z)=1, Scan range= 145 - 1450 m/z, Precursor Mass range= 380 - 980 m/z, ACG Target= 1000%, HCD collision energy= 25%).

Raw DIA data was analyzed with Spectronaut v19.0 (Biognosys) using the directDIA workflow while DDA data was analyzed with ProteomeDiscoverer™. MS2 data was used for searches against the *Anas platyrhynchos* UniprotKB database (https://www.uniprot.org). For method comparison and optimization, DDA was employed and search parameters were set as follows: precursor ion mass tolerance of 10 ppm and fragment ion mass tolerance of 0.02 Da. Trypsin was used as the digestion enzyme, allowing up to 2 missed cleavages. The false discovery rate (FDR) for peptide-spectrum match (PSM) was set to 1%. Proteins containing at least one unique peptide were considered for subsequent quantification and functional analysis.

### 2.7. Statistical analysis

Significant differences between the conditions tested were evaluated by one-way analysis of variance (ANOVA). Differences with *p* < 0.05 were considered statistically significant. Data were expressed as mean ± standard deviation.

## 3. Results and Discussion

### 3.1. Impact of Sample preparation strategy

#### 3.1.1. Experimental design and Protocol selection

The first step of the experimental design involved conducting a comprehensive bibliographic review to identify the most commonly used protocols for bottom-up proteomics sample preparation, particularly in the context of meat and cell cultivated products. Although similar steps are observed in most referenced protocols - protein solubilization, reduction and alkylation, digestion - (Phair et al., 2025), an overwhelming number of differences (reagents, time, temperature, materials) are observed, which may impact the effectiveness of protein preparation. These protocols with differences in time, workload, and cost can be grouped into 3 categories:

– Traditional in-solution digestion protocols: using chaotropic agents like urea or detergents such as Sodium dodecyl sulfate (SDS) or sodium deoxycholate (SDC). They are cost-effective (1–2 $/sample), highly reported in literature, but time-demanding as they involve multiple manual steps, which may not be compatible with the demanding fast pace needed in laboratories aiming for efficient repetition and scaling (Jiang et al., 2024).
– Device-based methods such as FASP, S-Trap, SP3, and commercial kits like PreOmics iST and EasyPep, which incorporate filtration or bead-based purification to improve cleanup and reproducibility (Varnavides et al., 2022). Despite being robust and fast (around 60 min of hand-on times), and therefore ideal for routine usage, these are considerably more expensive (30–50 $/sample) than conventional options, which understandably limits their adoption by several labs.
– Innovative in-solution sample prep approaches like Sample Preparation by Easy Extraction and Digestion (SPEED), which relies on acid-base extraction (Doellinger et al., 2020; Karousi et al., 2025). SPEED protocol is presented to offer advantages in terms of speed and cost-effectiveness. However, it has not yet been thoroughly validated for all proteomics applications, especially in cell-based meat.

#### 3.1.2. Comparison of sample preparation protocols

A comparison of five distinct protocols, representing three sample preparation categories, was conducted to assess their impact on the resulting protein profile (**Figure 1**).

**Figure 1.**
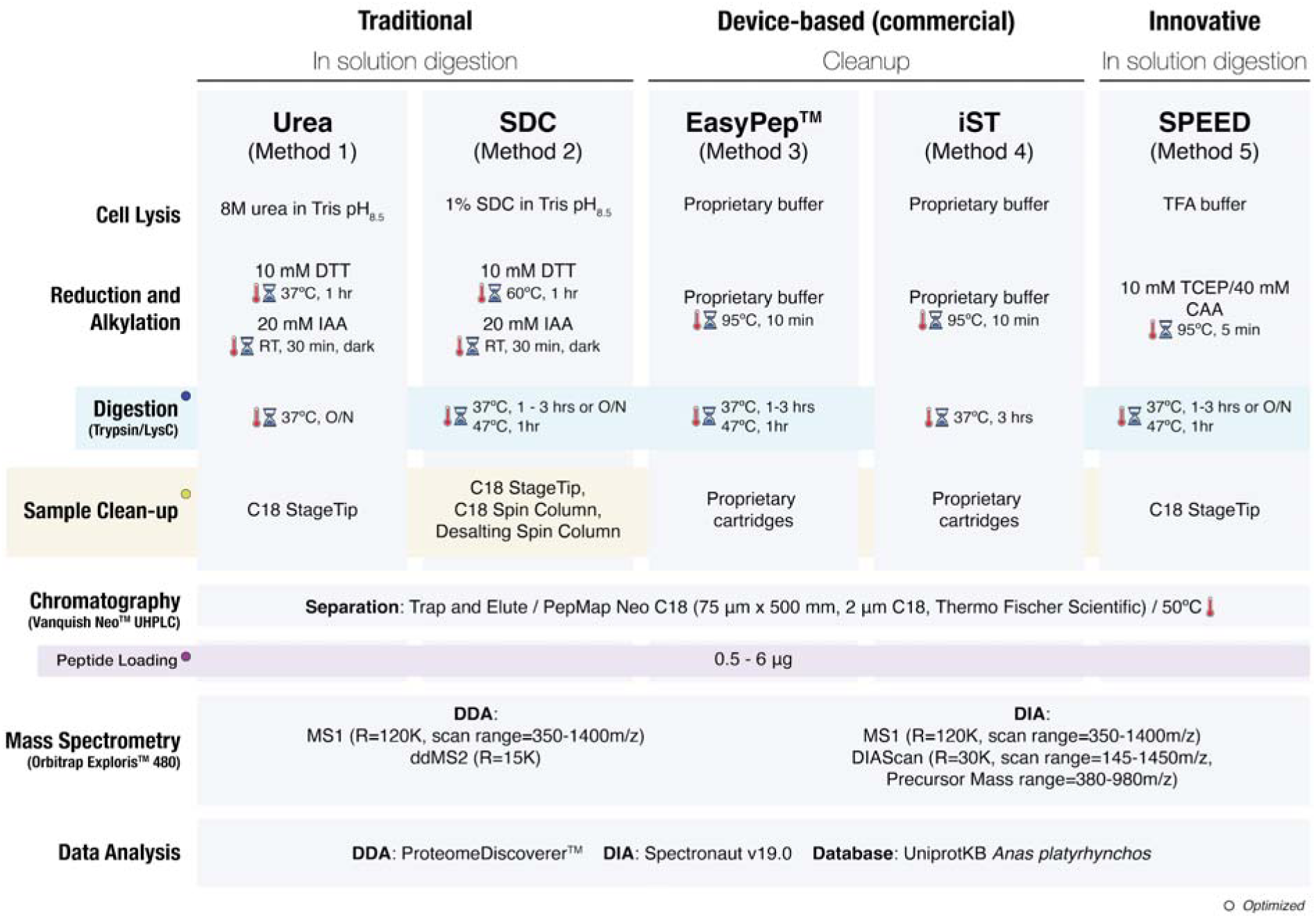
Experimental design: Comparison of five distinct protocols representing the three sample preparation categories reported in the literature and their different parameters and conditions.

##### Digested peptide yield comparison

Peptide quantification post-digestion revealed notable differences in yield across the five protocols tested. Overall, for 10µg of proteins extracted, device-based protocols provided the highest peptide yield. **Method 3** (136.4 ± 0.7 µg/mL) followed by **Method 4** (122.0 ± 4.4 µg/mL) consistently outperformed the tested in-solution methods. Among the latter, SPEED protocol (**Method 5:** 25.4 ± 1.0 µg/mL) led to higher peptide recovery than SDC (**Method 2:** 12.0 ± 1.1 µg/mL) and urea-based protocols (**Method 1:** 7.8 ± 2.9 µg/mL), likely due to improved protein solubilization and digestion under the tested conditions. These initial results highlighted the impact of sample preparation on peptide recovery, considering its importance in bottom-up proteomics results. As observed in **Figure 2 and Figure S1**, samples processed using device-based protocols (methods 3 and 4) showed a broader chromatographic range with peptides eluting at the earliest retention times (0 - 25 min). This suggests that device-based methods achieve more efficient digestion and recovery of smaller, hydrophilic peptides compared to conventional (urea, SDC) and SPEED protocols.

**Figure 2.**
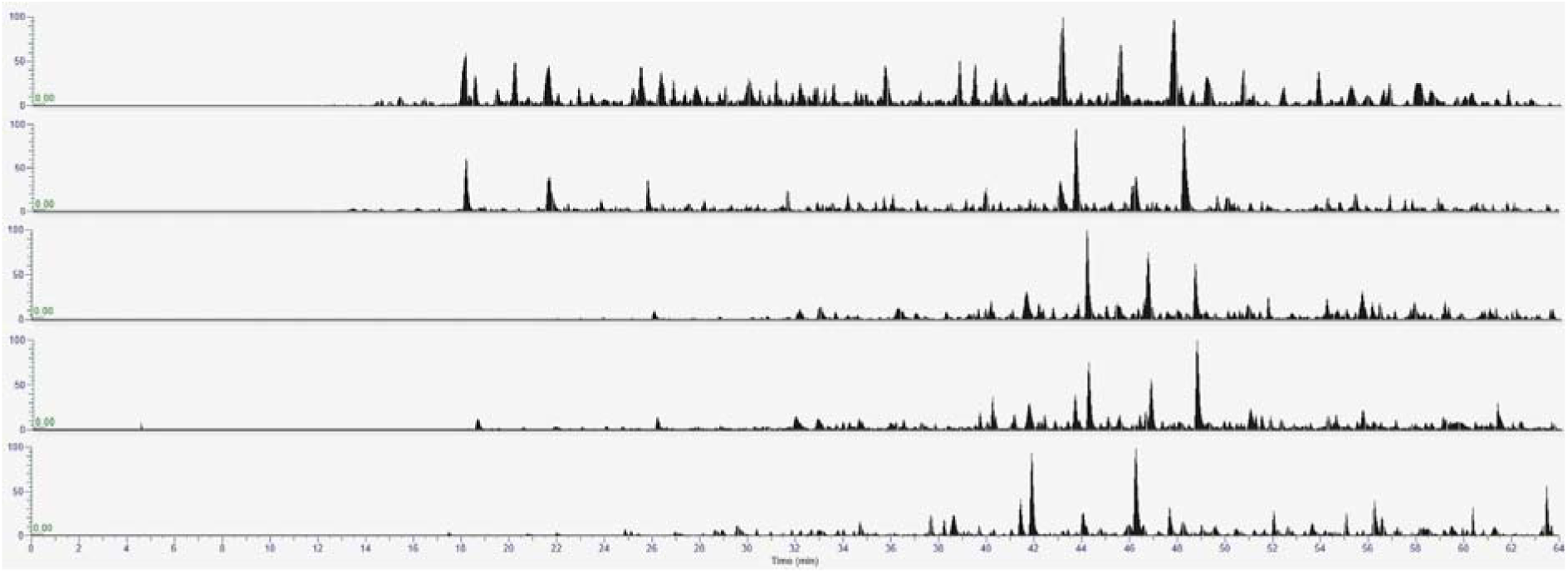
Overlaid base peak chromatograms of samples from different protocols. From top to bottom: Method 3 (Easypep kit), Method 4 (PreOmics kit), Method 2 (SDC based), Method 1 (Urea based) and Method 5 (SPEED).

##### Proteome coverage comparison across protocols

Protein identification results (**Figure 3**) further emphasized the performance differences among the five methods. Indeed, device-based protocols allowed the highest protein group identification highlighting their effectiveness in maximizing proteome coverage (**Figure 3B**). Notably, around 5100 proteins were identified with Method 3 followed by Method 4 (∼4150 IDs) and In-solution protocols (∼2500 - 3000 IDs). Peptide group identifications followed a similar trend to the protein IDs, with Method 3 yielding the highest IDs (∼29000) followed by iST (Method 4 ∼23850 IDs) and the other methods (∼14000 - 16000 IDs) (**Figure 3C**). Variations in the resulting protein profiles, as depicted in **Figure S2**, are also observed. Specifically, the heatmap illustrates distinct patterns of protein abundances across the various protocols.

**Figure 3.**
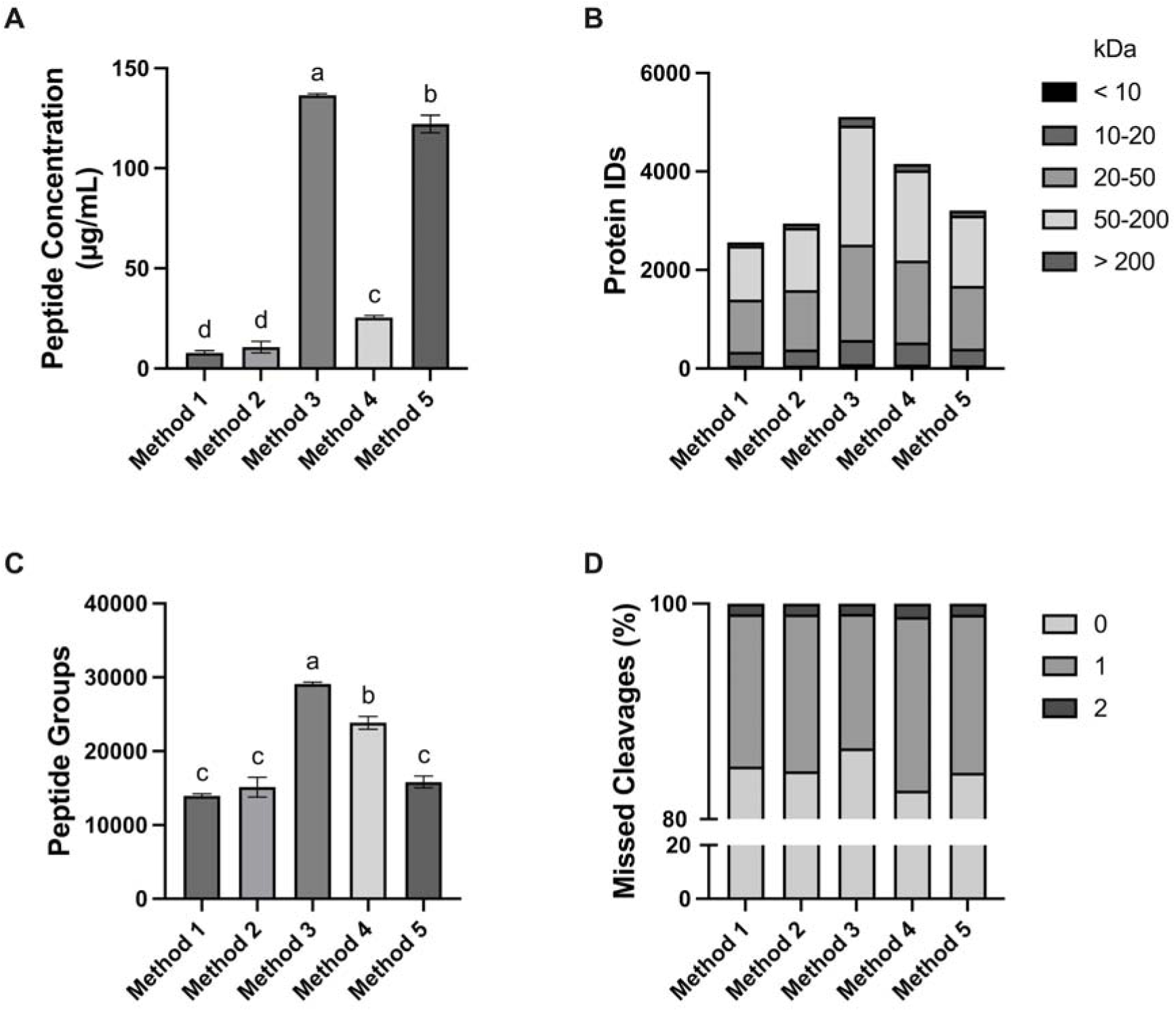
Comparison of proteomic results from different sample preparation methods. (A) Histogram showing a comparison of the peptide concentrations (µg/mL) for 10 µg of proteins extracted. (B) Total number of proteins identified with high confidence level and repartition by protein size (≤ 10KDa; 10-20 KDa; 20-50 KDa; 50 - 200 KDa; ≥ 200 KDa). (C) Total number of peptide groups identified with high confidence from each protocol. (D) Percentage of missed cleavages (0 - 2) between the identified peptide groups. Different lower case letters indicate statistically significant differences (*p* < 0.05).

Concerning the traditional in-solution protocols, although urea is the most reported detergent in bottom-up proteomics workflows, high urea concentrations can inhibit proteolytic enzymes through denaturation and may lead to protein carbamylation by ammonium cyanate, an urea degradation product. SDC, increasingly used in proteomics, is more LC-MS compatible and can boost trypsin activity, significantly improving protein digestion and sequence coverage (Danko et al., 2022). This could explain the tendency for higher protein IDs and peptide yield in Method 2 (SDC-based: ∼2950 IDs) compared to Method 1 (urea-based: ∼2550 IDs).

To further evaluate proteome coverage, the molecular weight (MW) distribution (**Figure 3B**) and gene ontology (GO) of the identified proteins was assessed (**Figure 4**). All protocols demonstrated broad coverage across a wide range of MW, from small proteins (<10 kDa) to large ones (>200 kDa) with EasyPep™ protocol (Method 3) showing the most comprehensive coverage. Indeed, through Method 3, the highest number of proteins in nearly every mass range was observed, particularly in 50–200 kDa categories, which typically represent the majority of cellular proteins. Similarly, Method 3 consistently yielded the highest number of protein IDs across all GO terms – Cellular Component, Biological Process, and Molecular Function – even though the relative distribution of the identified proteins remained consistent.

**Figure 4.**
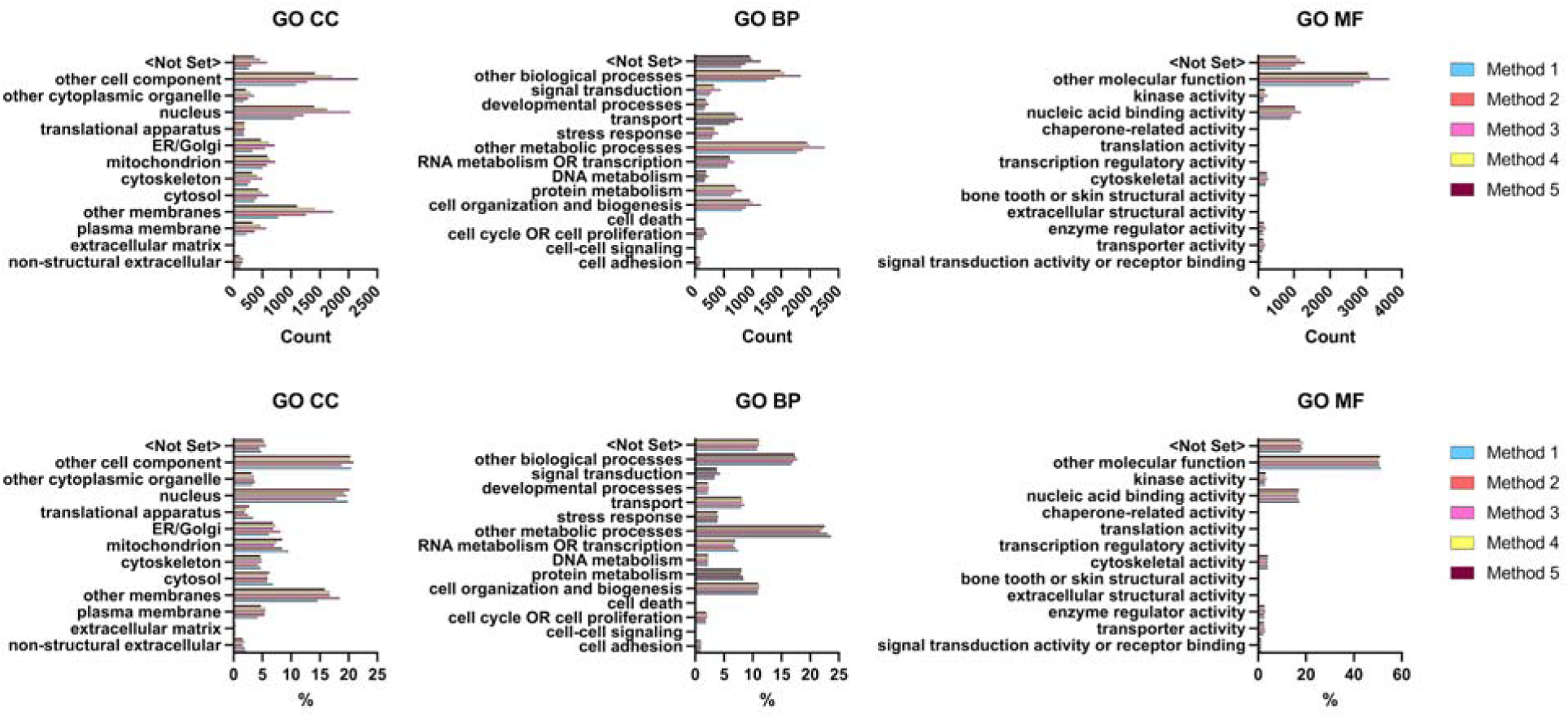
Comparison of the protein profile from different sample preparation methods through Gene Ontology (GO). GO analysis of the identified proteins is expressed in absolute counts (3 graphs on top) and in percentage of the total count (3 graphs of the bottom). The three categories of GO are represented: Cellular Component (CC), Biological Process (BP), and Molecular Function (MF). Method 1 (Urea based in-solution protocol); Method 2 (SDC based in-solution protocol); Method 3 (Easypep kit), Method 4 (PreOmics kit), and Method 5 (SPEED protocol).

To our knowledge, this study is the first to assess the impact of sample preparation on the proteome coverage of cultivated meat. Recently, Varnavides et al. (2022) compared 16 sample preparation methods (device-based and in-solutions approaches), which showed an overall similar performance on the proteome coverage of HeLa cells. In that study, SDC-based protocols achieved protein identification levels comparable to high-performing device-based methods (iST, FASP, EasyPep) and slightly surpassed other in-solution approaches (SPEED, urea-based).

The superior performance of the device-based protocols in this study may be attributed to their high optimization and standardization, using proprietary reagents and cleanup strategies. This likely improves protein solubilization, digestion efficiency, and peptide recovery, while better preserving hydrophilic and low-abundance peptides recovery. Nevertheless, the difference between protocols indicates that with further optimization, in-solution protocols may approach device-based performance levels.

### 3.2. Effect of digestion conditions

From the first comparison, methods 2, 3, and 5 were selected to further assess the impact of digestion conditions on the proteome results. The efficiency of the digestion conditions was measured in terms of digested peptides yield (µg/mL) and protein identifications.

#### 3.2.1. Device-based method digestion optimization (Method 3 EasyPep)

To determine the optimal digestion conditions for the EasyPep™ protocol, four conditions were tested: 1, 2, and 3 h at 37 °C and 1 h at 47 °C. Notably, the tested conditions were based on manufacturer user guide, recommending a digestion time between 1 to 3 hours. Although the proteome coverage remained stable (RSD ≤ 1%) across the tested conditions (**Figure 5D**), the yield of digested peptides exhibited a slight downward trend over time with 1 h at 37°C yielding 75.2 ± 2.0 µg/mL, closely followed by 1 h at 47°C (72.6 ± 0.8 µg/mL), 2 h (70.6 ± 3.1 µg/mL) and 3 h (65.6 ± 2.8 µg/mL) at 37°C (**Figure 5A**). The findings demonstrate that a short digestion time, such as 1h at 37 °C, is optimal for sample preparation performance. Furthermore, the consistent proteomics results observed across different conditions confirm the high robustness of this device-assisted protocol for cultivated meat application.

**Figure 5.**
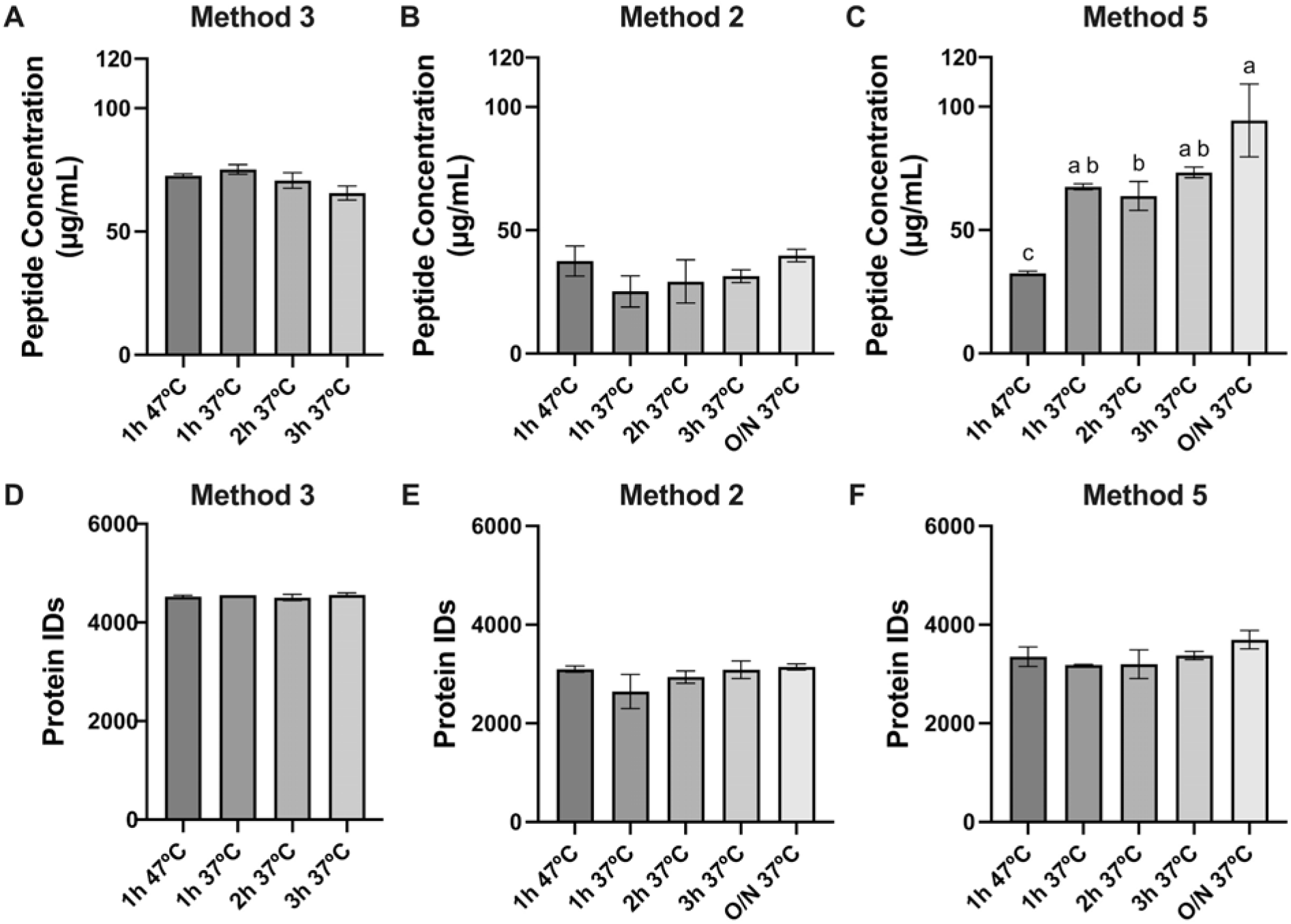
Effect of digestion conditions (1 h at 47 °C, 1h at 37 °C, 2h at 37 °C, 3h at 37 °C, and overnight at 37 °C) on peptide concentration (µg/mL) and total protein identification. Peptide quantification results for each digestion condition expressed in µg/mL in Method 3 (A. EasyPep™ based protocol), Method 2 (B. SDC-based protocol) and Method 5 (C. SPEED protocol). D, E, and F: Total number of proteins identified across digestion conditions in Method 3, 2, and 5 respectively. O/N overnight. Values are expressed as mean ± standard deviation. Different lower case letters indicate statistically significant differences (*p* < 0.05).

#### 3.2.2. In-solution digestion optimization (Method 2)

To assess the impact of digestion conditions for the SDC-based protocol, five conditions were tested: 1 h at 47 °C (a), 1h at 37 °C (b), 2h at 37 °C (c), 3h at 37 °C (d), and overnight at 37 °C (e). For 10 µg of proteins digested, peptide yield was 37.5 ± 6.1 µg/mL, 25.2 ± 6.3 µg/mL, 29.3 ± 8.8 µg/mL, 31.4 ± 2.6 µg/mL, and 39.8 ± 2.6 µg/mL, for conditions a, b, c, d, and e, respectively (**Figure 5B**). In 37°C conditions, protein identifications also showed a tendency to slightly increase with time, yielding 3021 IDs at 1h, 3243 IDs at 2h, 3391 IDs at 3h, and 3420 IDs overnight (**Figure 5E**). Notably, digestion at 47 °C produced a comparable peptide yield and protein coverage (3372 IDs after 1h at 47°C) to the 3h and overnight condition, suggesting some flexibility in protocol timing.

#### 3.2.3. Method 5 (SPEED) digestion optimization

SPEED digestion was tested under the same five conditions compared for Method 2. 47°C digestion resulted in the lowest peptide yield (32.4 ± 0.9 µg/mL), while 37°C digestions yield 67.5 ± 1.2 µg/mL; 63.8 ± 5.8 µg/mL; 73.4 ± 2.2 µg/mL; and 94.5 ± 14.8 µg/mL, for conditions b, c, d, and e respectively. However, protein IDs results were as follows: 3350 (a), 3182 (b), 3197 (c), 3373 (d), and 3697 (e), showing a slight upward trend with digestion time at 37°C (**Figure 5F**). Doellinger et al. (2020) introduced the SPEED as a fast and superior alternative to detergent/chaotropic agent-based protocols. SPEED was presented as a method to simplify and standardize sample preparation, while enhancing proteome coverage, particularly in challenging samples. Although the authors recommended an overnight (37 °C) digestion period, the effect of digestion time on the SPEED results was not investigated.

Proteome analysis using a heat map clustering (**Figure 6**) revealed that overnight digestion favored a specific subset of high-abundance proteins, and a specific profile which was different from the reduced digestion time conditions (1 - 3h). Notably, the 3h digestion showed a more balanced expression, capturing features from both ends of this spectrum and achieving a broad and even proteome representation. Taken together, 3h digestion at 37 °C can be considered optimal for both SDC and SPEED protocols, offering a good balance between comprehensive data and efficient processing, avoiding the drawbacks of long overnight incubations.

**Figure 6.**
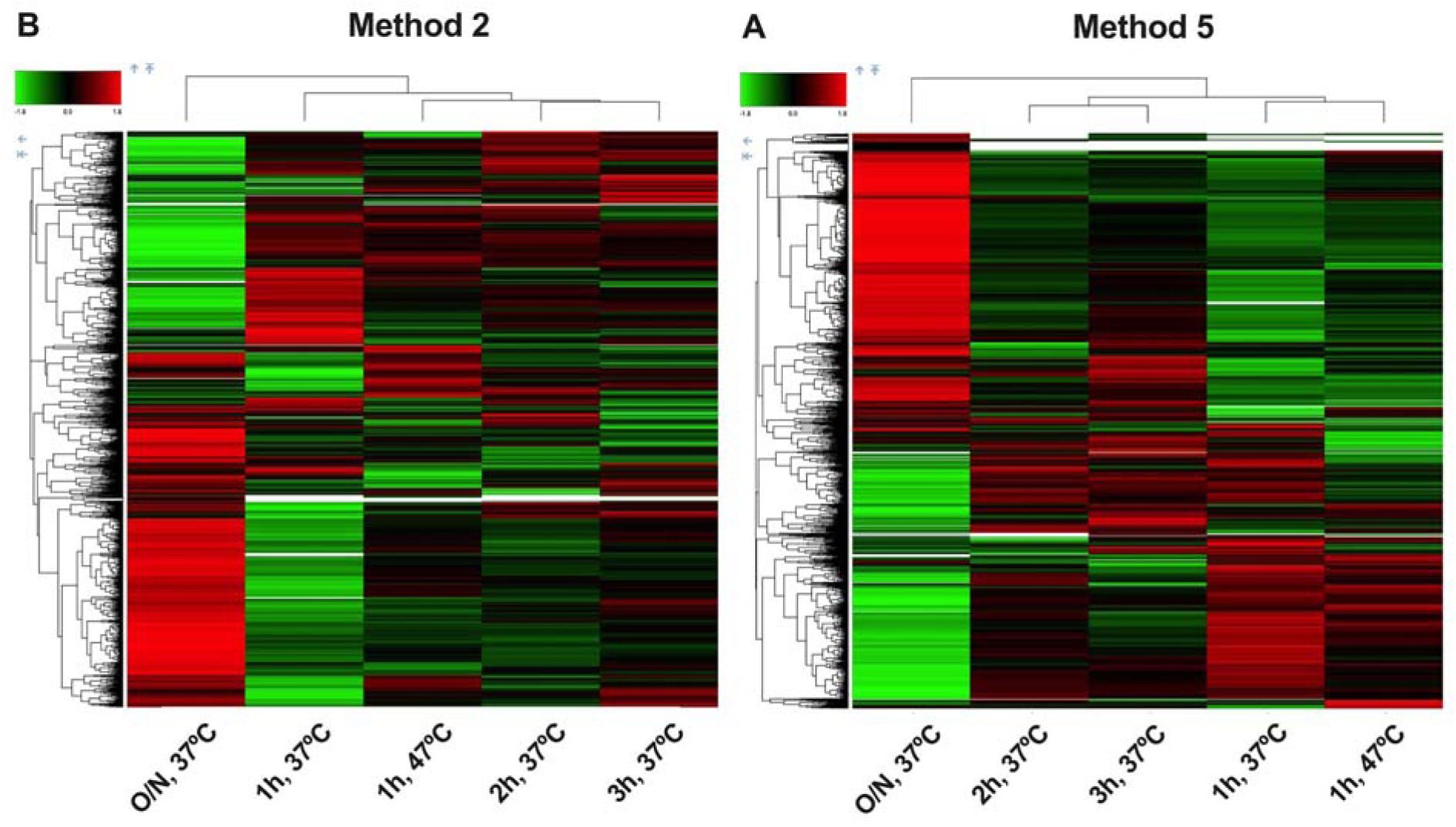
Heatmap, hierarchical clustering of normalized protein abundances (grouped) across different digestion conditions: 1h 47°C; 1h 37°C; 2h 37°C; 3h 37°C; and Overnight 37°C. From left to right: Heatmap of Method 2 (SDC protocol) and Method 5 (SPEED). The lines in the heatmap represent the relative abundance of proteins across the conditions. On the upper left side of the figures is a scale indicating the color code relative to the normalized protein abundance (ranging from −2.8 to 2.8). Dendrogram depicts hierarchical relationships of clusters based on euclidean distance function and complete linkage method.

### 3.3. Impact of Peptide cleanup strategy

After digestion, peptide cleanup prior to ESI-MS analysis is crucial for the removal of salts and reagents that can interfere with the ionization process (matrix effect, spectral complexity) and reduce the lifespan of the analytical system (Tubaon et al., 2017). To evaluate the impact of peptide cleanup on sample recovery and proteome depth, three commercial SPE devices were tested after SDC-based protocol: C18 StageTips, Pierce™C18 Spin Columns (centrifuge column with C18 resin), and Pierce™ Desalting Spin Columns (made with a proprietary polymer-based hydrophobic resin).

Peptide groups and protein identifications varied depending on the cleanup approach. The proteome coverage shows a higher yield of peptide groups and protein identifications by the Pierce™ Peptide Desalting Column cleanup (∼ 26700 peptide groups / 4380 protein IDs), outperforming C18 cleanup (19000 - 21000 peptide groups / 3840 - 3960 IDs) (**Figure 7**). Additionally, the amino acid distribution profile remains unchanged regardless of the clean-up technology used; only the total number of protein identifications increases proportionally. Common desalting is based on C18 solid phase extraction (SPE) tips, as peptides are retained to C18 material while salts elute in aqueous acidic environments. However, the desalting is associated with peptides lost, which impact the peptide recovery and potentially the final proteome profile due to inadequate retention or irreversible adsorption onto C18 materials (Kanao et al., 2024). These results emphasize the importance of cleanup strategies within the specific context of cultivated meat proteomics as sample preparation performance can vary significantly with the cleanup workflow applied.

**Figure 7.**
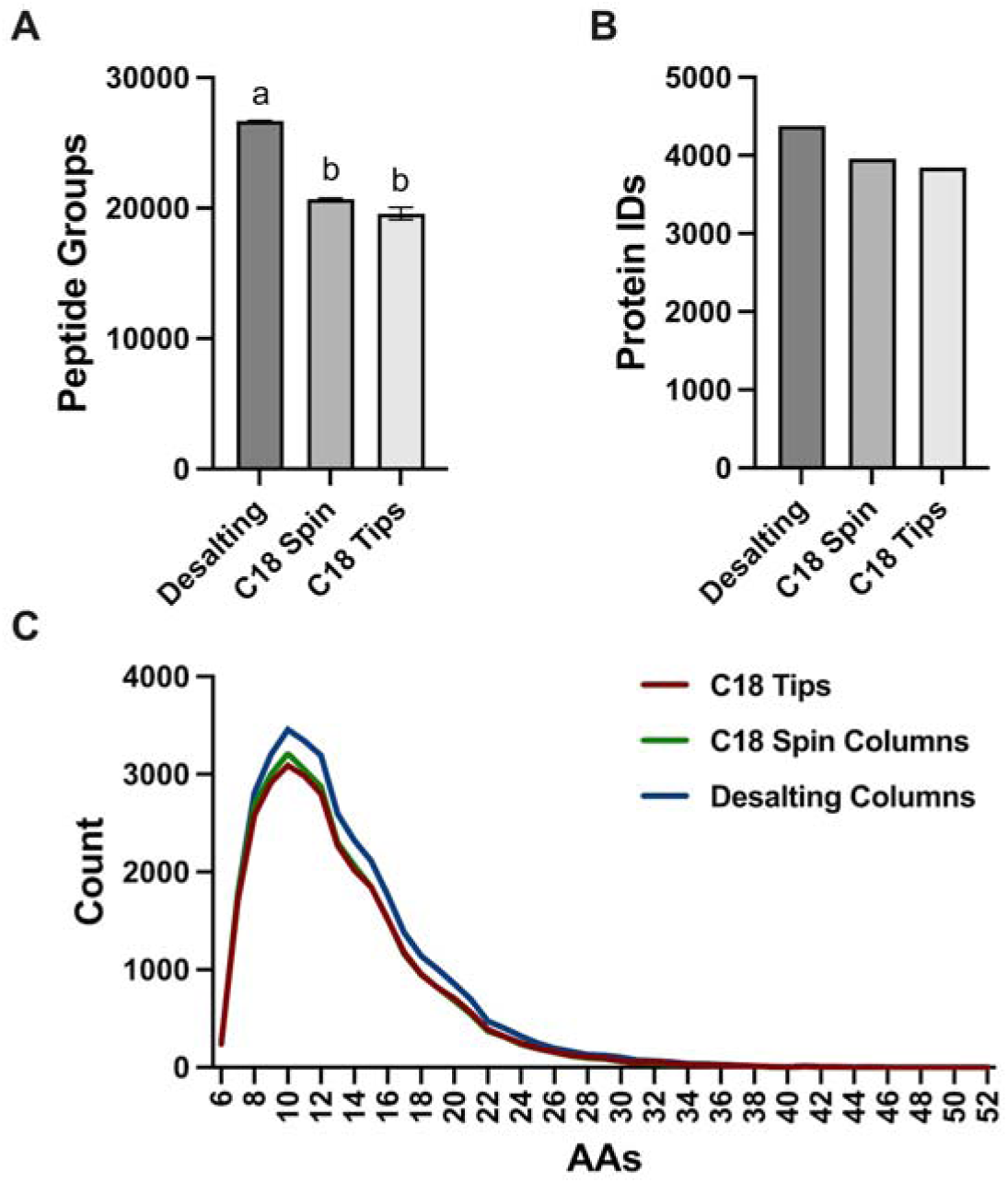
Impact of peptide cleanup strategy. **(A)** Total number of peptide groups identified with high confidence from each cleanup method. **(B)** Total number of proteins identified across clean-up conditions. **(C)** Amino acids (AAs) distribution of peptides recovered from different clean-up conditions. Different lower case letters indicate statistically significant differences (*p* < 0.05).

### 3.4. Impact of LC-MS workflow

#### Mass spectrometry acquisition workflow

Data-Dependent Acquisition (DDA) and Data-Independent Acquisition (DIA) are the two widely used strategies in MS-proteomics. In this study, both workflows were tested and the results obtained were compared (**Figure S3**). DIA provided a more thorough proteome coverage compared to DDA, identifying about 20% more protein groups (around 6,500 vs 5,000 IDs) with higher precision (CV% of protein abundances: 7.6% vs 12.8% for DIA and DDA respectively). The difference in performance between workflows can be explained by their differences in data acquisition approach. Indeed, DDA first performs a mass spectrometry scan (MS1) to gather m/z and abundance data, then selects and fragments specific precursor ions (MS2) for peptide identification. The stochastic selection of precursor ions can limit the detection of low-abundance ions leading to a reduced run-to-run reproducibility. DIA overcomes this by eliminating precursor selection, instead systematically fragmenting all ions within fixed or dynamic isolation windows, which results in improved data completeness and reproducibility with less missing values (Ghosh et al., 2025). With advances in data analysis algorithms, DIA has shown superior identification efficiency compared to DDA, leading to its growing application in various research fields (Zou et al., 2025). However, with its longer development history, DDA benefits from a more established software and database ecosystem, widely adopted by analytical platforms.

#### Peptide loading

Moreover, the impact of the peptide loading amount (0.5 – 6 ug) was also assessed to investigate the optimal peptide amount to be injected (**Figure 8**). Injection of 500 ng only revealed a fraction of the proteome indicating that injection of an increased amount can boost peptide and thus protein identifications. Moreover, the abundance profile of proteins varies with the injection amount (**Figure S4**). Analysis suggests 2 μg as an optimal peptide amount to be loaded on LC-MS to increase peptide identifications while limiting overload of nanoLC columns.

**Figure 8.**
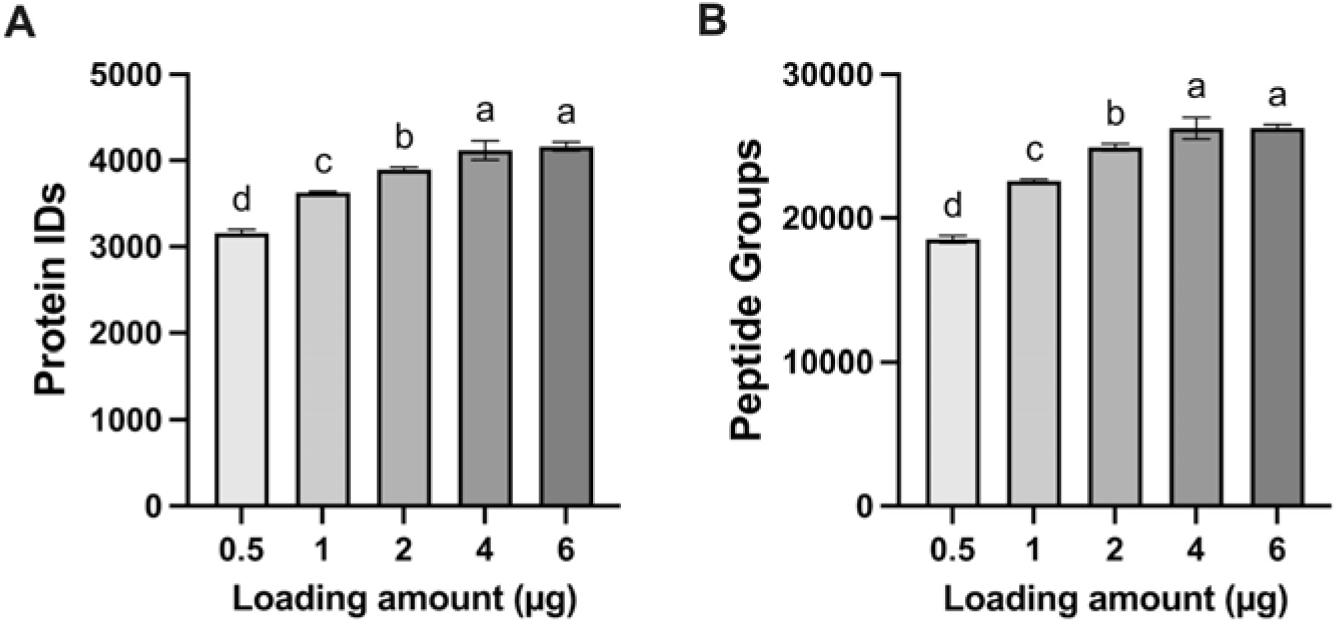
Impact of peptide loading amount (0.5 - 6µg) on the proteome coverage. From left to right, histograms illustrate the impact of loading amount on the identification of peptides and proteins. Different lower case letters indicate statistically significant differences (*p* < 0.05).

### 3.5. Discussions

#### 3.5.1. Opportunities and Challenges of proteomics adoption in the cultivated meat industry

##### Research & development opportunities

As cultivated meat production is fundamentally associated with providing alternative protein sources, the comprehensive characterization of its protein profile has a significant added value. Indeed, combining proteomics with classic protein quantification methods such as the Kjeldahl method and total amino acid analysis, provides a deeper understanding of the protein profile, thereby efficiently supporting multiple research and development challenges faced by the industry. For example, the assessment of the scaling effect and processes variation on protein quality and stability, or the demonstration of nutritional parity with conventional meat can be supported by comparing the meat proteome at different conditions (Eldrid et al., 2025). Moreover, a multi-omics integration of transcriptomic, proteomic, and metabolomic layers, can provide a system-level understanding of mechanisms linked to cell differentiation, cell growth or media adaptation (Chen et al., 2025; Trautmann et al., 2025). Proteomics could play a future role in cultivated meat quality and authentication assessment, defining specific protein/peptide markers that can indicate the quality and origin of the cultivated products (Sidira et al., 2024; Agregan et al., 2023).

##### Safety assessment opportunities

With the growing demand for NAMs as animal-free methods to assess the safety of novel foods such as cultivated meat, multi-omics is increasingly taken into consideration. For example, multi-omics approaches can help to verify the absence of unintended protein overexpression, genetic drift and confirm the safety profile established by foundational omics layers (Deng et al., 2025). Transcriptomics stand out as the most advanced and widely applied omics approach for novel food hazard characterization due its standardized workflows (high robustness) and deepened coverage. Proteomics can serve as a useful complement to transcriptomics, providing functional verification of the phenotype by detailing protein structure and epitope configuration (Ham et al., 2025). Moreover, proteomics could serve to verify the absence of protein residues used in the culture media (e.g. growth factors) and assess the impact of food processing (heating, cooking, etc) on putative allergen stability. This approach is already used in the conventional meat industry where MS-proteomics is employed to study meat allergens (Carrera et al., 2024). Recently, Yang et al. (2025) proposed an LC-MS/MS method for the quantification of livestock and poultry meat allergens. Integrating proteomic data with genomic and/or transcriptomic layers can enhance the accuracy of these *in silico* models, making them more comparable to *in vivo* results. This multi-layered evidence directly supports the transition toward the full replacement of animal procedures, in line with legal directives and frameworks governing scientific safety assessments (Directive 2010/63/EU).

##### Proteomics adoption challenges

The global adoption of proteomics in the cultivated meat industry is challenging due to the inherent method complexity and the lack of standardized guidelines for the generation of proteomic data in a regulatory context. The high technical complexity associated with proteomics, particularly in applications such as PTM (glycosylation or phosphorylation), top-down or single cell analysis (Dowling et al., 2023; Guo et al., 2025) represents a barrier for their routine use despite their promising added-value for the cultivated meat industry. Even shot-gun proteomics, while more and more adopted owing to its simplified applications and the supporting ecosystem (preparation device, data acquisition workflows and processing software) still face the issue of confidence in protein annotation, which is particularly crucial in cultivated meat for drawing valid conclusions. In bottom-up proteomics, protein annotation is fundamentally an iterative process of sequence alignment and statistical inference. Following the acquisition of MS/MS spectra, raw data is processed through search engines (e.g., Mascot, MaxQuant, or SEQUEST) to perform Peptide-Spectrum Matching (PSM). These experimental spectra are correlated against theoretical fragmentation patterns derived from a defined FASTA protein database (Jiang et al., 2024). The selection of this database is a critical parameter in the experimental design, as it dictates the balance between annotation depth and False Discovery Rate (FDR). If Swiss-Prot, the reviewed section of UniProtKB is the gold standard for high-confidence annotation as every entry is manually curated by expert biologists, not all animal species benefit from a deep Swiss-Prot coverage. Therefore, scientists are obliged to use unreviewed databases (TrEMBL or NCBI RefSeq), which are wider but also associated with higher redundancy. Furthermore, the data processing pipeline used (algorithms, softwares, FDR control, etc) can also significantly impact the protein profile result (Lou et al., 2023; Frankenfield et al., 2025).

Interlaboratory or method variations like sample preparation conditions, data acquisition setups and processing methods can influence the proteomic profiles of cultivated meat, posing a challenge to data interpretation in safety assessment. For example, *In-silico* allergenicity prediction, which compares novel food protein sequences to known allergens (sliding 80-amino acid window: >35% similarity, >50% identity), may yield different conclusions depending on proteome coverage. As observed in this study, the protein profile obtained varied according to the sample preparation strategy (device-based, in solution, SPEED protocol), the cleanup strategy, or LC-MS acquisition workflows (DDA or DIA) with up to 45% difference in protein group identification (**Figure 3**). This high disparity directly impacts data fidelity and carries the risk of non-detection and restricted proteome coverage, with potential impact on analytical conclusions.

#### 3.5.2. Standardization of proteomics workflows for cultivated meat

To overcome these challenges and ensure reliable and comparable data, the standardization of proteomic methods is essential. This includes the implementation of certified workflows and stringent quality control (QC) measures. Shotgun proteomics presents the advantage of being widely adopted by analytical platforms, and therefore can represent a starting base for a standardized application of proteomics in the regulatory assessment of cultivated meat.

Concerning the sample preparation standardization, the results obtained suggest that device-based protocols with optimized parameters (e.g. Easy pep protocol, 37°C of digestion for 1h) provide a deep proteome coverage with high robustness allowing reliable data comparison between different platforms. The optimized SPEED and SDC-based protocols (supplementary materials) can represent a cost-effective alternative with comparable efficiency and throughput to device-based methods. Regarding digestion conditions, a critical parameter in bottom-up proteomics, our results identify 3-hours at 37 °C as the optimal balance between workflow efficiency and analytical output for cultivated meat proteomics. Furthermore, polymer-based hydrophobic resin (Pierce™ Peptide Desalting Spin Columns), consistently yielded high peptide recovery and superior protein identification levels. **Figure 9** shows a global comparison of 03 sample preparation protocols – Easypep, SDC, and SPEED – before and after optimization. In terms of protein IDs, the optimized in-solution protocols (∼4500 - 5000 IDs) demonstrated performance highly comparable to the Easypep method. The protocol optimization substantially decreased the differences in protein group exclusivity from 30% (1500 protein groups unique to Easypep method before optimization) to 10%. Despite this, differences in protein abundances and overall profiles remain apparent across the various methods, as illustrated by the heatmap clustering (**Figure 10**). Consequently, proteomic data derived from different cultivated meat samples or conditions cannot be statistically compared if varied sample preparation techniques were utilized.

**Figure 9.**
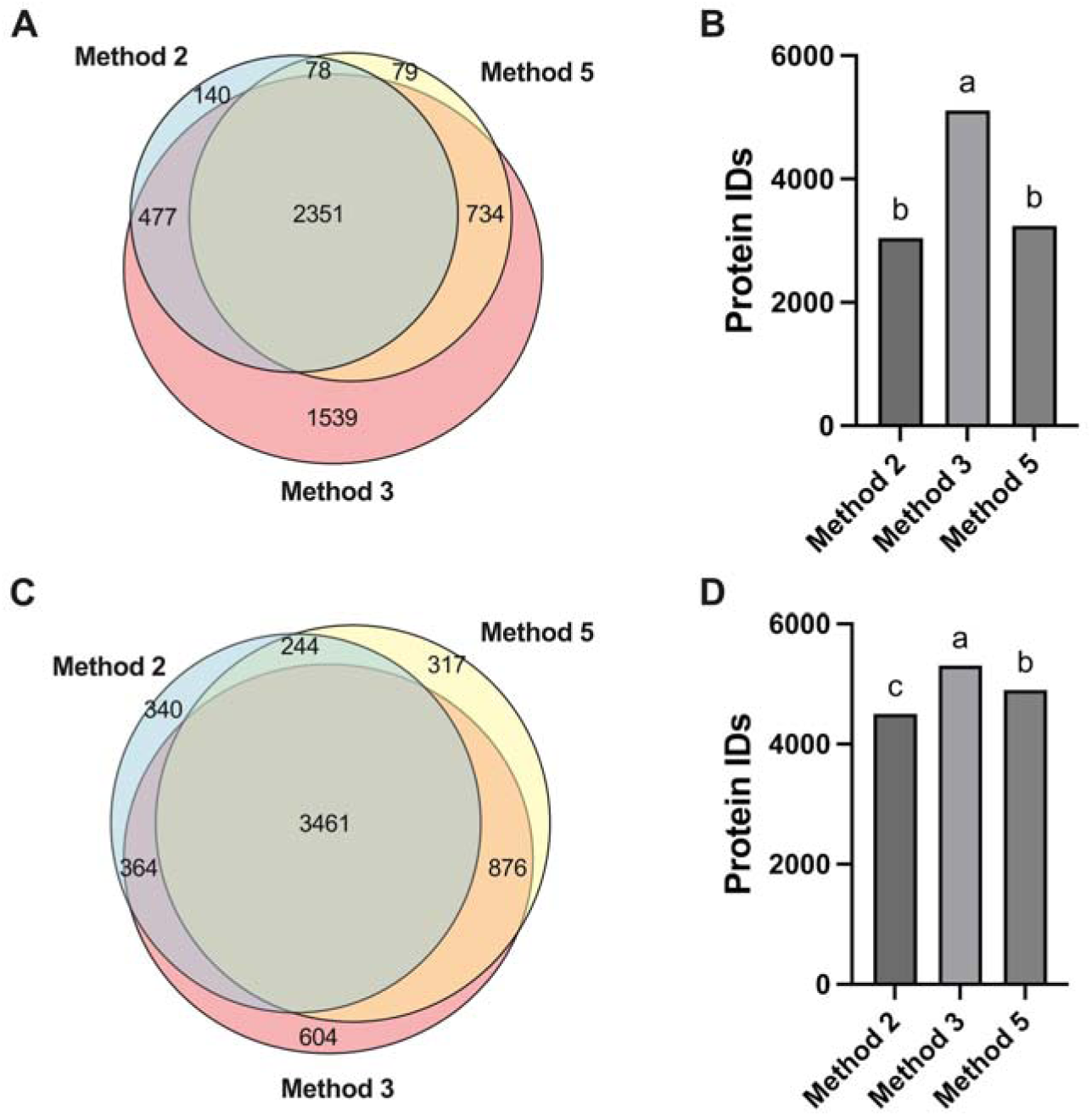
Comparison of proteome coverage across sample preparation protocols – Method 2 (SDC-based protocol), Method 3 (EasyPep™ based protocol), and Method 5 (SPEED protocol) – before (on top) and after (bottom) optimization. Venn diagrams showing the total number of proteins differentially expressed across the protocols. Histograms showing the total number of proteins identified across protocols. Different lower case letters indicate statistically significant differences (*p* < 0.05).

**Figure 10.**
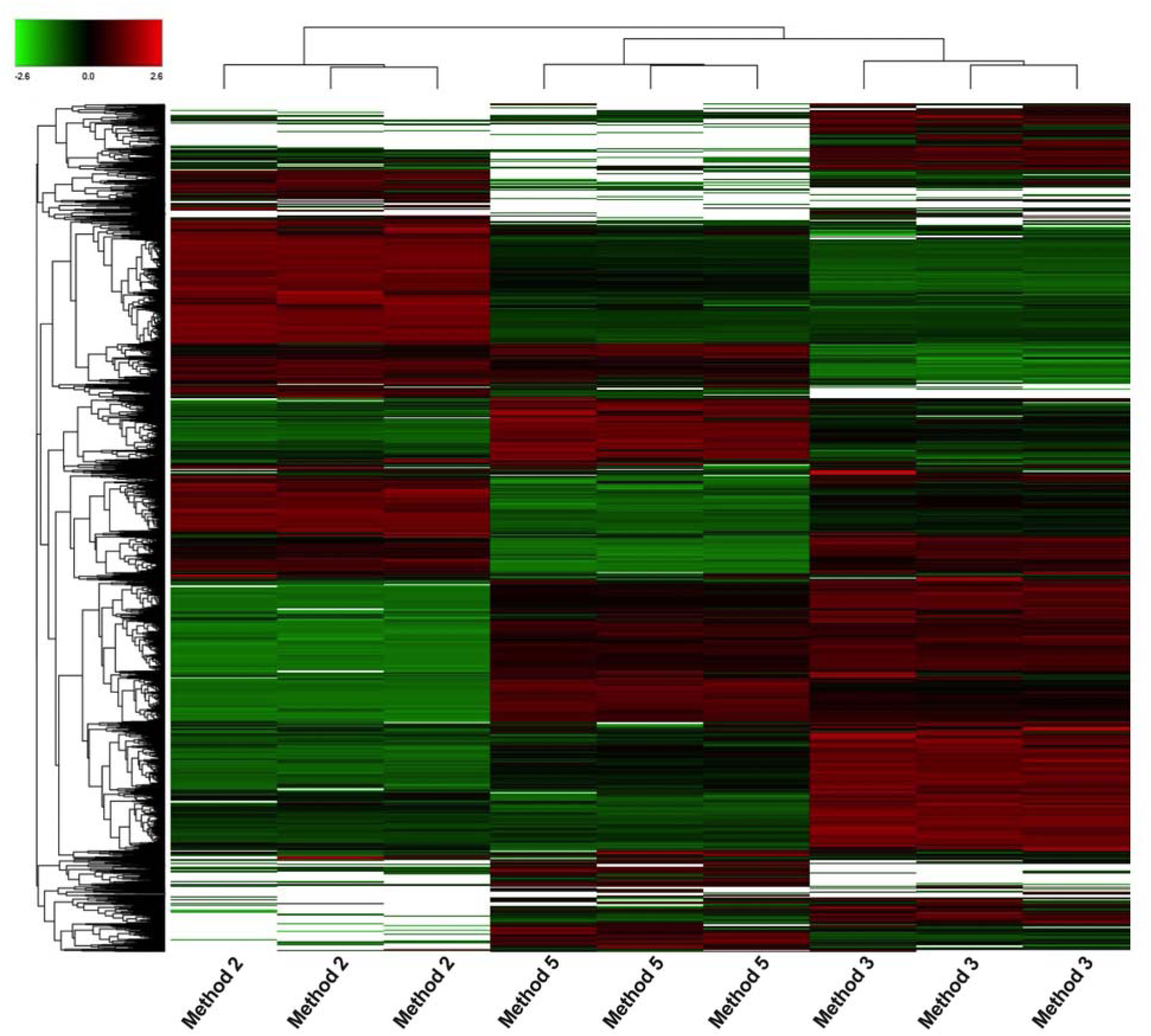
Comparison of optimized protocols. Heatmap, hierarchical clustering of normalized protein abundances across samples (*n*=3) from different optimized protocols: Method 2 (SDC-based protocol), Method 3 (EasyPep™ based protocol), and Method 5 (SPEED protocol). The lines in the heatmap represent the relative abundance of proteins across the optimized conditions. On the upper left side of the figures is a scale indicating the color code relative to the normalized protein abundance (ranging from −2.6 to 2.6). Dendrogram depicts hierarchical relationships of clusters based on euclidean distance function and complete linkage method.

Variations in proteomics can also arise from the MS acquisition workflow. Indeed, while DDA is the most used workflow in proteomic application, DIA, as shown in this study, offers superior identification and deeper proteome coverage. However, DDA still faces utility in specific applications such as multiplexing or PTM proteomics. For the latter, DDA offers superior localization potential, which is crucial for identifying specific peptides and accurately mapping complex PTM sites (Zou et al., 2025). Conversely, DIA yields more identifications but makes it more challenging to pinpoint the exact localization site of modifications, such as phosphorylation. Furthermore, DDA is currently the main viable workflow for multiplexed quantification strategies like tandem mass tag (TMT)-based quantification. This multiplexing approach is highly efficient, allowing for the concurrent analysis of multiple experimental conditions in a single LC-MS injection. The benefits include maximized instrument efficiency, reduced analytical time (from days to hours), fewer missing values, and higher accuracy in relative quantification (Kang et al., 2026). Therefore, the ultimate choice between DDA and DIA must be made based on a comprehensive evaluation of the study goals and requirements.

#### 3.5.3. Guidelines for proteomics application in cultivated meat studies

Proteomics data has the potential to effectively complement existing transcriptomics-based approaches for novel food characterization. For instance, the Food Standards Agency (FSA) already recommends applicants to provide the qualitative and quantitative data on the macro- and micronutrient composition of their Cell Cultivated Products (CCP). This requires also defining the protein profile of the CCP, including any post-translational modifications where relevant, and comparing it to the conventional counterpart (FSA, 2025). However, consideration has not yet been given to the strategy and analytical workflows that must be used for such proteomics studies. This study presents validated and cost-effective Standard Operating Procedures (SOPs), specifically the SPEED and SDC-based methods, for bulk bottom-up proteomics sample preparation, tailored for cultivated meat applications (details in Supplementary materials).

Examining the application of proteomics in cultivated meat studies, we offer the following recommendations to guide authorities and stakeholders in this context:

– Proteomics as a complementary layer in the weight of evidence approach: Proteomics should be considered as complementary data within a broader weight of evidence approach. For example, to evaluate the comparative nutritional profiles, traditional assays such as total amino acid analysis and in vitro digestibility can be effectively complemented by shotgun proteomics to benchmark the protein profiles of cultivated meat products against their conventional analogues, provided that both meat products are analyzed through the same workflow. Indeed, the use of similar workflows for cultivated meat and their conventional counterparts is fundamental in reducing the variability inherent to shotgun proteomics analysis.
– Methodological Transparency and Validation: Transparency in the methodology (sample preparation, data acquisition and processing) should be required, including the full disclosure in the instrument used, the search engine, software version, etc. Moreover, protein annotation should prioritize curated databases or supported by precise documentation of FASTA file origins and data processing pipelines, when curated databases are poor or not available.
– Integrated Allergenicity Assessment: When assessing allergenicity, assessors should note that sample preparation and LC-MS methods can significantly influence the proteome coverage. This limitation may hinder the effective screening of potential allergens associated with the undetermined proteins. Therefore, proteomics should not be used in isolation or replace other omics strategies such as transcriptomics or genomics but integrated into a multi-omics strategy. Such a framework can be used to facilitate comprehensive cell-culture meat safety assessment as suggested by Mathieu et al. (2025).
– Harmonization of Analytical Standards: Regulators and industry stakeholders should collaborate in order to recognize accepted protocols (sample preparation, LC-MS, and data processing workflows) and narrow analytical variability. A framework should be established for the acceptance of cost-effective sample preparation protocols (such as optimized SDC) or data processing software (DIA-NN) as viable alternatives to high-cost device-based methods and proprietary platforms, provided that applicants can demonstrate equivalent performance in terms of proteome coverage and data quality. Moreover, regulators should strongly encourage the inclusion of standardized QC such as Hela digest in all proteomic submissions to ensure interlaboratory data comparability.

## Conclusions

In this study, we rigorously addressed for the first time the critical lack of standardization in proteomics for cultivated meat by systematically evaluating the influence of key bottom-up workflow variables on the proteome profile of cultivated meat. This standardization will be essential for generating reliable data necessary for multi-omics integration (especially complementing transcriptomics in NAMs) and ultimately supporting robust quality assessment, R&D efforts, and future regulatory submissions in the novel food sector.

## Acknowledgements

We thank Brian Chun and Hannah Lester for the regulatory review of the manuscript, and the whole team of PARIMA for fruitful discussion and providing cultivated duck biomass.

## Declaration of generative AI and AI-assisted technologies in the manuscript preparation process

During the preparation of this work the authors used “Gemini” in order to correct English and enhance the readability of the paper. After using this tool, the authors reviewed and edited the content as needed and assume full responsibility over the content of the published article.

## Author contributions: CRediT

João Palma: Investigation, Software, Data curation, Visualization, Writing – original draft.

Chloé Colchero Leblanc: Investigation, Methodology, Visualization

Remy Kusters: Writing – review and editing, Resources

Franks Kamgang Nzekoue: Conceptualization, Supervision, Writing – review and editing

## Reference list

Agregan, R., Pateiro, M., Kumar, M., Franco, D., Capanoglu, E., Dhama, K., & Lorenzo, J. M. (2023). The potential of proteomics in the study of processed meat products. Journal of Proteomics, 270, 104744. 10.1016/j.jprot.2022.104744

Bakhsh, A., Kim, B., Ishamri, I., Choi, S., Li, X., Li, Q., … & Park, S. (2025). Cell-Based Meat Safety and Regulatory Approaches: A Comprehensive Review. Food science of animal resources, 45(1), 145. doi: 10.5851/kosfa.2024.e122

Carrera, M., Abril, A. G., Pazos, M., Calo-Mata, P., Villa, T. G., & Barros-Velázquez, J. (2024). Proteins and peptides: proteomics approaches for food authentication and allergen profiling. Current Opinion in Food Science, 57, 101172. 10.1016/j.cofs.2024.101172

Chen, N., Niu, Q., Ye, Q., Wu, Q., Wu, Y., Zhao, X., … & Chen, L. (2025). Accelerating cultivated meat bioprocess innovation: Rational design of cost-effective serum-free media via the convergence of data-driven analytics and synthetic biology platforms. Trends in Food Science & Technology, 105526. 10.1016/j.tifs.2025.105526

Danko, K., Lukasheva, E., Zhukov, V. A., Zgoda, V., & Frolov, A. (2022). Detergent-assisted protein digestion—on the way to avoid the key bottleneck of shotgun bottom-up proteomics. International Journal of Molecular Sciences, 23(22), 13903. 10.3390/ijms232213903

Deng, M., Zhang, Z., Ahmed, I., Yong, L., Li, Q., Lin, H., & Li, Z. (2025). A Review on Allergens, Allergenicity Assessment Methods, and Specific Allergenicity Assessment Strategies for Novel Foods. Journal of Agricultural and Food Chemistry, 73(43), 27177–27188. 10.1021/acs.jafc.5c08212

Directive, E. (2010). 63/EU of the European Parliament and of the Council of 22 September 2010 on the protection of animals used for scientific purposes. Off. J. Eur. Union, 276, 33–79. https://eur-lex.europa.eu/LexUriServ/LexUriServ.do?uri=OJ:L:2010:276:0033:0079:en:PDF

Doellinger, J., Schneider, A., Hoeller, M., & Lasch, P. (2020). Sample preparation by easy extraction and digestion (speed) - a universal, Rapid, and detergent-free protocol for proteomics based on Acid Extraction. Molecular & Cellular Proteomics, 19(1), 209–222. 10.1074/mcp.tir119.001616

Dowling, P., Swandulla, D., & Ohlendieck, K. (2023). Mass spectrometry-based proteomic technology and its application to study skeletal muscle cell biology. Cells, 12(21), 2560. 10.3390/cells12212560

EFSA Panel on Dietetic Products, Nutrition and Allergies (NDA), Turck, D., Bresson, J. L., Burlingame, B., Dean, T., Fairweather-Tait, S., … & van Loveren, H. (2016). Guidance on the preparation and presentation of an application for authorisation of a novel food in the context of Regulation (EU) 2015/2283. Efsa Journal, 14(11), e04594. 10.2903/j.efsa.2016.4594

Eldrid, C., Hawke, E., Cain, K. M., Meeson, K., Watson, J., Spiess, R., … & Barran, P. (2025). From Discovery to Delivery: A Rapid and Targeted Proteomics Workflow for Monitoring Chinese Hamster Ovary Biomanufacturing. Molecular & Cellular Proteomics, 101011. 10.1016/j.mcpro.2025.101011

Food Standards Agency. (2025). Supplementary guidance to applicants for assessment of cell cultivated products (CCP) in food: Allergenicity & nutrition. Retrieved from https://www.food.gov.uk/regulated-products/supplementary-guidance-to-applicants-for-assessment-of-cell-cultivated-products-ccp-in-food-allergenicity-nutrition / Accessed on 04 December 2025

Frankenfield, A. M., Yang, K. L., binti Mazli, W. N. A., Shih, J., Yu, F., Lo, E., … & Hao, L. (2025). Benchmarking SILAC Proteomics Workflows and Data Analysis Platforms. Molecular & Cellular Proteomics, 24(6). 10.1016/j.mcpro.2025.100980

Ghosh, G., Shannon, A. E., & Searle, B. C. (2025). Data acquisition approaches for single cell proteomics. Proteomics, 25(1-2), 2400022. 10.1002/pmic.202400022

Gu, H., Kong, Y., Huang, D., Wang, Y., Raghavan, V., & Wang, J. (2025). Scaling cultured meat: challenges and solutions for affordable mass production. Comprehensive reviews in food science and food safety, 24(4), e70221. 10.1111/1541-4337.70221

Guo, T., Steen, J. A., & Mann, M. (2025). Mass-spectrometry-based proteomics: from single cells to clinical applications. Nature, 638(8052), 901–911.10.1038/s41586-025-08584-0

Ham, J. H., Lee, Y. J., Lee, I., & Kim, H. Y. (2025). Allergenicity in cultured meat: assessment and strategic management. Critical Reviews in Food Science and Nutrition, 1–13. 10.1080/10408398.2025.2497919

Hu, M., & Wang, Y. (2024). Optimized workflow for proteomics and phosphoproteomics with limited tissue samples. Current Protocols, 4(4). 10.1002/cpz1.1028

Jiang, Y., Rex, D. A. B., Schuster, D., Neely, B. A., Rosano, G. L., Volkmar, N., … & Meyer, J. G. (2024). Comprehensive overview of bottom-up proteomics using mass spectrometry. ACS Measurement Science Au, 4(4), 338–417. 10.1021/acsmeasuresciau.3c00068

Kanao, E., Tanaka, S., Tomioka, A., Ogata, K., Tanigawa, T., Kubo, T., & Ishihama, Y. (2024). High-Recovery Desalting Tip Columns for a Wide Variety of Peptides in Mass Spectrometry-Based Proteomics. Analytical Chemistry, 96(52), 20390–20397. 10.1021/acs.analchem.4c03753

Kang, C., Hong, J., Kim, H., Jo, J., Seo, J. H., Lee, J. W., & Lee, S. W. (2026). A Robust Strategy for High-Throughput and Deep Proteomics by Combining Narrow-Window Data-Independent Acquisition and Isobaric Mass Tagging. Journal of Proteome Research. 10.1021/acs.jproteome.5c00501

Kardas, M., Staśkiewicz-Bartecka, W., & Kołodziejczyk, A. (2025). Cultured Meat Reformulation: Health Potential and Sustainable Food Challenges—Narrative Review. Comprehensive Reviews in Food Science and Food Safety, 24(6), e70262. 10.1111/1541-4337.70262

Karnaneedi, S., Limviphuvadh, V., Maurer-Stroh, S., & Lopata, A. L. (2023). De Novo Transcriptomic Analyses to Identify and Compare Allergens in Foods. In Food Allergens: Methods and Protocols (pp. 351–365). New York, NY: Springer US. 10.1007/978-1-0716-3453-0_24

Karousi, P., Voumvouraki, M., Nikolaou, P. E., Kollias, I., Paradeisi, F., Sampanai, E., … & Courraud, J. (2025). Easy Proteomics Sample Preparation: Technical Repeatability and Workflow Optimization Across 8 Biological Matrices in a New Core Facility Setting. Proteomics, 25(20), 15–24. 10.1002/pmic.

Kaulich, P. T., & Tholey, A. (2025). Top-Down Proteomics: Why and When?. Proteomics, 25(24), 6–12. 10.1002/pmic.202400338

Kim, S., Jeong, Y., Jo, H., Park, Y. G., & Moon, S. H. (2025). Cultured meat: advances in stem cell biology, tissue engineering, and bioprocess optimisation for scalable and sustainable production—a review. International Journal of Food Science and Technology, 60(2), vvaf220. 10.1093/ijfood/vvaf220

Ledwith, R., Stobernack, T., Bergert, A., Bahl, A., Pink, M., Haase, A., & Dumit, V. I. (2024). Towards characterization of cell culture conditions for reliable proteomic analysis: in vitro studies on A549, differentiated THP-1, and NR8383 cell lines. Archives of Toxicology, 98(12), 4021–4031. 10.1007/s00204-024-03858-4

Lee D. Y., Hur, S. J. (2025). Gaps and solutions for large scale production of cultured meat: a review on last findings. Current Opinion in Food Science, 61, 101243. 10.1016/j.cofs.2024.

Levine, S. L., Riter, L. S., Lagadic, L., Bejarano, A. C., Burden, N., Burgoon, L. D., … & Armbrust, K. L. (2025). Challenges and Opportunities in the Development and Adoption of New Approach Methods (NAMs). Journal of Agricultural and Food Chemistry, 73(13), 7519–7521. 10.1093/toxres/tfae044

Lou, R., Cao, Y., Li, S., Lang, X., Li, Y., Zhang, Y., & Shui, W. (2023). Benchmarking commonly used software suites and analysis workflows for DIA proteomics and phosphoproteomics. Nature Communications, 14(1), 94. 10.1038/s41467-022-35740-1

Mathieu, T., Légaré, S., Nzekoue, A. F., Jauré, N., Lester, H., Dias, T., & Kusters, R. (2025). Integrative multi-omics modelling for cultivated meat production, quality, and safety. Trends in Food Science & Technology, 105364. 10.1016/j.tifs.2025.105364

Molho, D., Ding, J., Tang, W., Li, Z., Wen, H., Wang, Y., … & Tang, J. (2024). Deep learning in single-cell analysis. ACM Transactions on Intelligent Systems and Technology, 15(3), 1–62. dl.acm.org/doi/10.1145/3641284

Phair, I. R., Sovakova, M., Alqurashi, N., Nisr, R. B., McNeilly, A. D., Lamont, D., & Rena, G. (2025). In-depth proteomic profiling identifies potentiation of the LPS response by 7-ketocholesterol. Journal of Molecular and Cellular Cardiology Plus, 11, 100285. 10.1016/j.jmccpl.2025.100285

Prabahar, A., Zamora, R., Barclay, D., Yin, J., Ramamoorthy, M., Bagheri, A., … & Jiang, P. (2024). Unraveling the complex relationship between mRNA and protein abundances: a machine learning-based approach for imputing protein levels from RNA-seq data. NAR Genomics and Bioinformatics, 6(1), lqae019. 10.1093/nargab/lqae019

Qiu, S., Kratochvilova, E., Huang, W. E., Cui, Z., Agnew, T., Yang, A., & Ye, H. (2026). Proteome constrained metabolic modeling of Sus scrofa muscle stem cells for cultured meat production. Metabolic Engineering. 10.1016/j.foodres.2025.116627

Schar, S., Rass, L., Malinovska, L., Savickas, S., Cavallo, F., Below, C., … & Reiter, L. (2025). A flexible end-to-end automated sample preparation workflow enables standardized and scalable bottom-up proteomics. Analytical Chemistry, 97(40), 22116–22131. 10.1021/acs.analchem.5c03829

Sidira, M., Smaoui, S., & Varzakas, T. (2024). Recent proteomics, metabolomics and lipidomics approaches in meat safety, processing and quality analysis. Applied Sciences, 14(12), 5147. 10.3390/app14125147

Singh, S., Yadav, R., Hussain, S., & Singh, A. N. (2026). Cultured meat: Current status, challenges, and strategic prospects. In One Planet, One Health, One Future (pp. 495-506). Academic Press. 10.1016/B978-0-443-38325-0.00032-0

Trautmann, C. L., Ghosh, A., Kalkan, A. K., Noé, F., & Bar-Nur, O. (2025). Enhanced Media Optimize Bovine Myogenesis in 2D and 3D Models for Cultivated Meat Applications. Advanced Science. DOI: 10.1002/advs.202413998

Tubaon, R. M., Haddad, P. R., & Quirino, J. P. (2017). Sample clean-up strategies for ESI mass spectrometry applications in bottom-up proteomics: Trends from 2012 to 2016. Proteomics, 17(20), 1700011. DOI: 10.1002/pmic.201700011

Varnavides, G., Madern, M., Anrather, D., Hartl, N., Reiter, W., & Hartl, M. (2022). In search of a universal method: a comparative survey of bottom-up proteomics sample preparation methods. Journal of proteome research, 21(10), 2397–2411. 10.1021/acs.jproteome.2c00265

Woodland, B., Farrell, L. A., Brockbals, L., Rezcallah, M., Brennan, A., Sunnucks, E. J., … & Padula, M. P. (2025). Sample Preparation for Multi-Omics Analysis: Considerations and Guidance for Identifying the Ideal Workflow. *Proteomics*, e13983. 10.1002/pmic.13983

Yang, S., Shenchen, Y., Zhao, Y., Zhou, M., Abdallah, M. F., Gao, Y., … & Li, Y. (2025). Development and validation of an LC-MS/MS method for a sensitive quantitation of livestock and poultry meat allergens. Food Chemistry, 145965. 10.1016/j.foodchem.2025.145965

Zou, X., Wang, L., Chen, Y., Fu, H., Gao, Y., Liu, B., … & Zhai, L. (2025). In-depth analysis of data characteristics and comparative evaluation of dda and dia accuracy in label-free quantitative proteomics of biological samples. Clinical Proteomics. 10.1186/s12014-025-09572-2

