## Supplementary materials for "Proteomics for cultivated meat: the importance of Analytical Standardization"

**D**

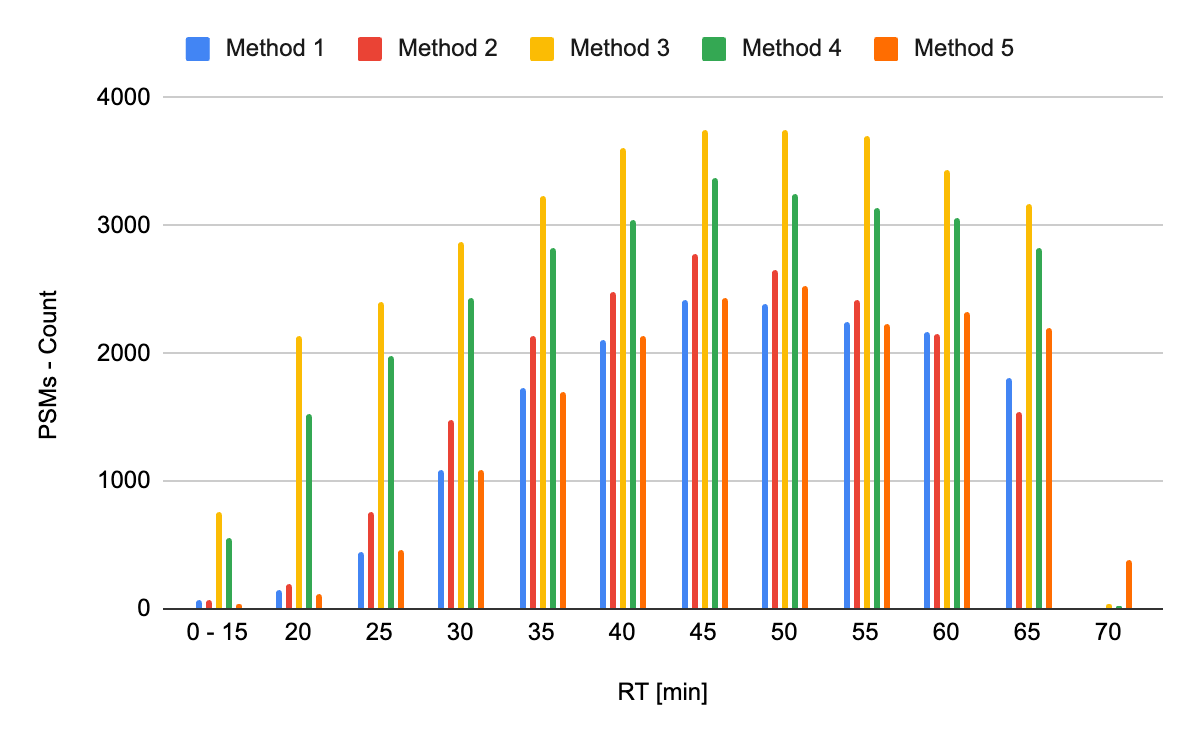

**Figure S1.** Distribution of peptide identifications along retention time (RT) for each sample preparation method. PSM: peptide spectrum matches

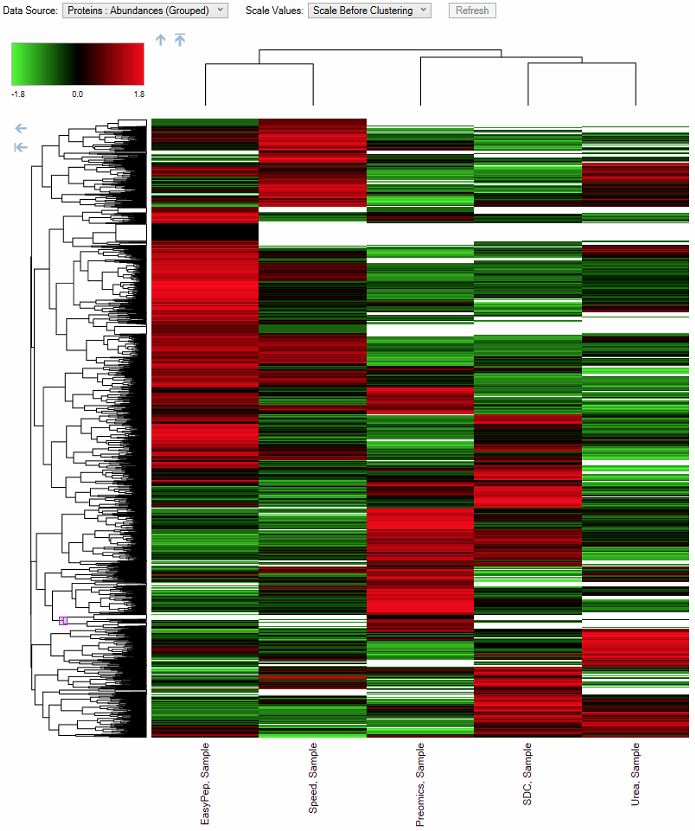

**Figure S2.** Heatmap, hierarchical clustering of normalized protein abundances (grouped) from compared sample preparation protocols (*n*=3): (from left to right) EasyPep (Method 3); SPEED (Method 5); Preomics (Method 4); SDC (Method 2); Urea (Method 1). The lines in the heatmap represent the relative abundance of proteins across the sample prep. On the upper left side of the figure is a scale indicating the color code relative to the normalized protein abundance (ranging from -1.8 to 1.8). Dendrogram depicts hierarchical relationships of clusters based on euclidean distance function and complete linkage method.

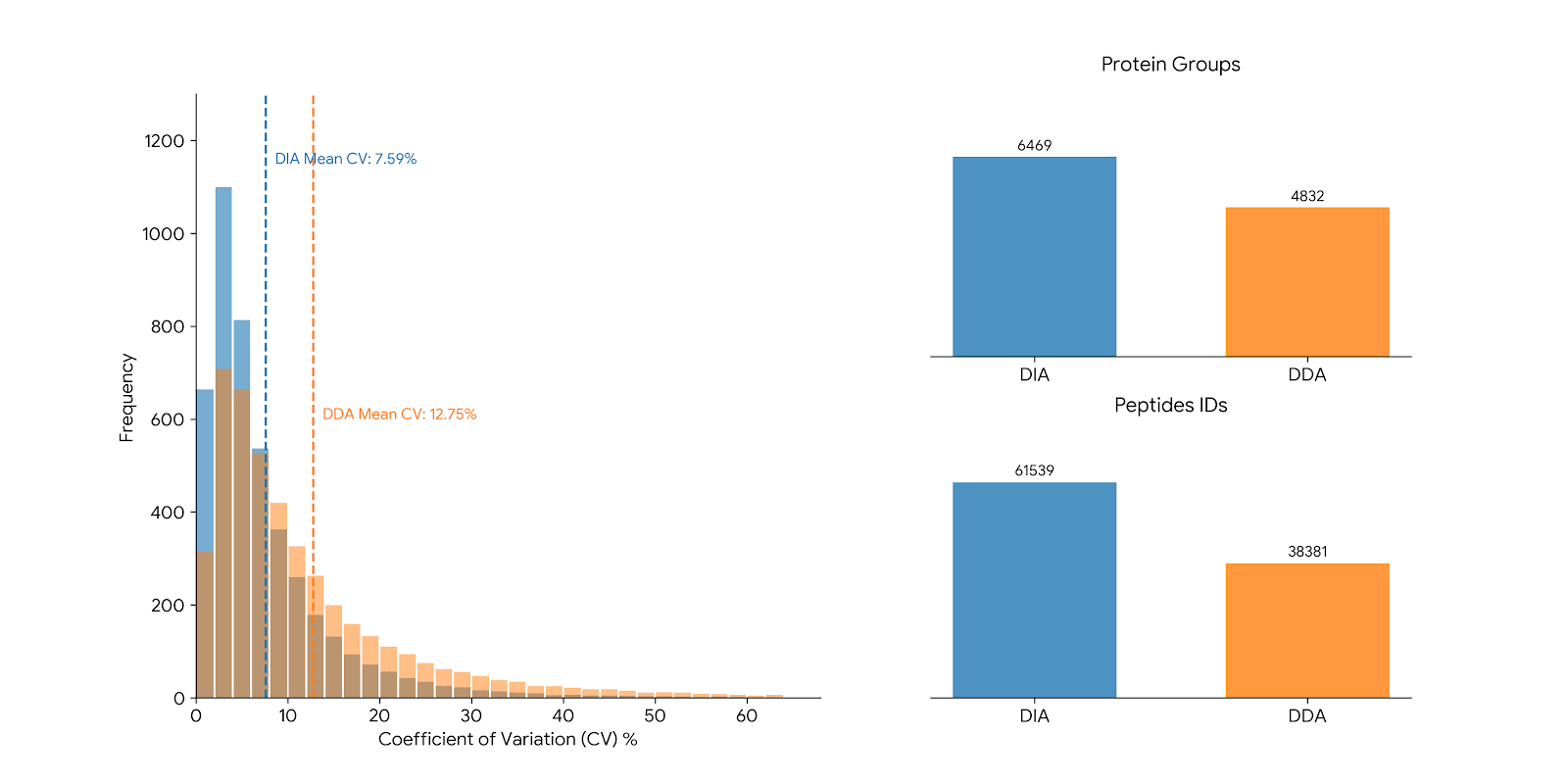

**Figure S3.** Data dependent acquisition (DDA) and Data Independent acquisition (DIA) performance comparison for cultivated meat proteomics (precision, protein and peptide IDs). CV coefficient of variation (%) of abundances of the identified proteins. DDA and DIA results are in orange and blue, respectively.

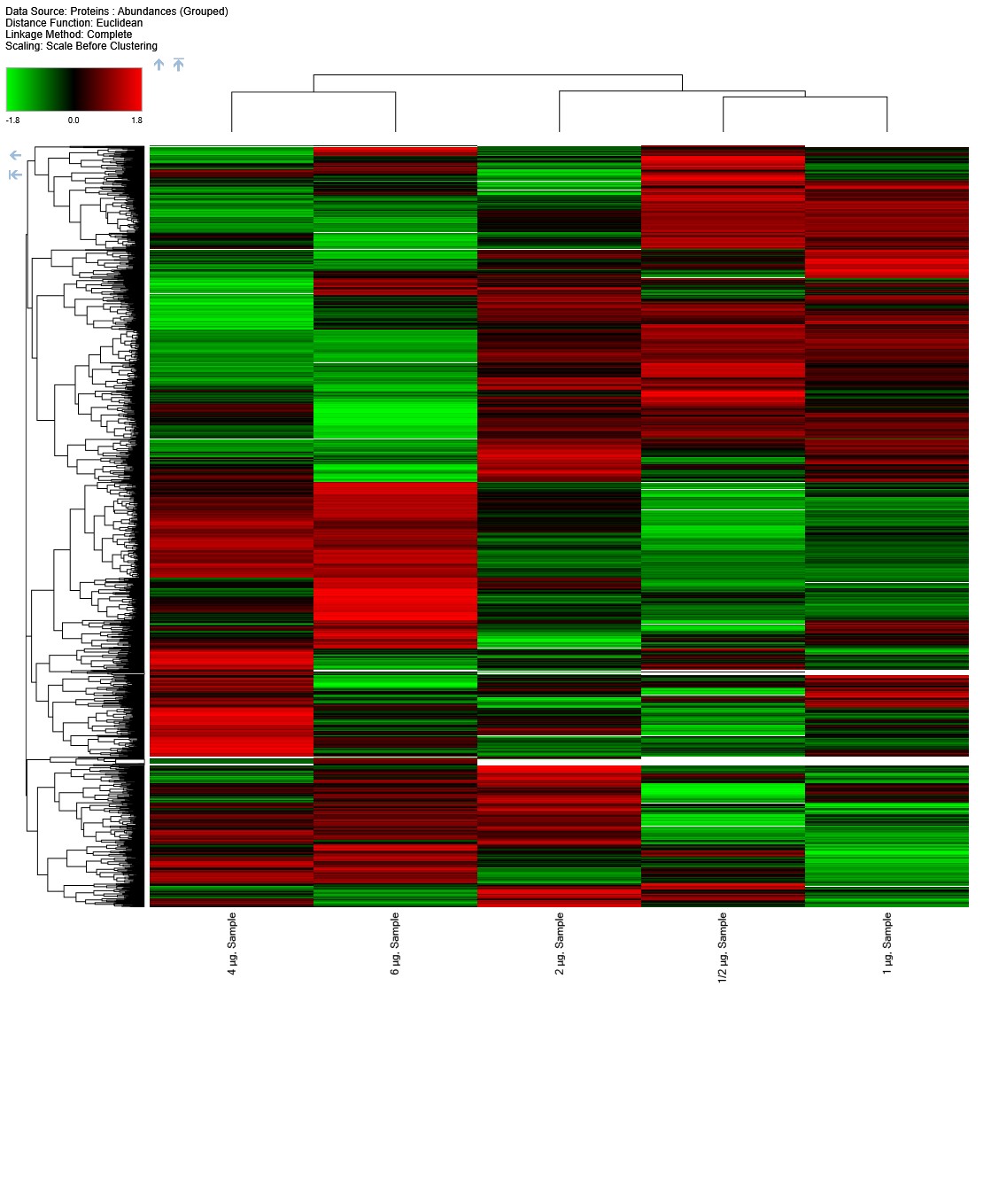

**Figure S4.** Heatmap, hierarchical clustering of normalized protein abundances (grouped) from compared peptide loading amount (*n*=3). From left to right: 4 µg; 6 µg; 2 µg; 0.5 µg; 1 µg. The lines in the heatmap represent the relative abundance of proteins across the sample injections. On the upper left side of the figure is a scale indicating the color code relative to the normalized protein abundance (ranging from -1.8 to 1.8). Dendrogram depicts hierarchical relationships of clusters based on euclidean distance function and complete linkage method.

**Bottom-up proteomics – Cost-effective sample-preparation protocols:**

A. **Sodium deoxycholate (SDC) - in solution protocol**

**1. Protein Lysis, Reduction, and Alkylation**[1](https://docs.google.com/document/d/1nYrCoQfv9UJzQDybHZKV2XnNJie5t592iQ-DkVvmbII/edit)

1. Resuspend the cell pellet in Lysis Buffer (1% SDC in 0.1 M Tris-HCl, pH 8.5) until the sample is homogenous. Consider a ratio of 1 mg pellet to 5-10µL of lysis buffer (ideally this ratio should be optimized for each cell type).
2. Incubate the sample for 10 minutes at 60°C in a ThermoMixer set to 500 rpm.
3. Centrifuge the samples at 16,000 rcf for 20 minutes at 4°C.
4. Carefully collect the supernatant containing the solubilized proteins and transfer it to a new tube.
5. A protein concentration assay can be performed at this point to adapt the protein content. We suggest having 100 to 200 µg of protein content for the next step.
6. Perform a reduction step by adding 50 µL of 10 mM Dithiothreitol (DTT). Vortex and incubate for 1 hour at 60°C (0 rpm).
7. Alkylate the sample by adding 100 µL of 20 mM Iodoacetamide (IAA). Vortex.
8. Incubate the sample in the dark for 30 minutes at room temperature (RT).

**2. Proteolytic Digestion**

9. Add Trypsin/Lys-C protease mix based on a 1:20 enzyme-to-protein ratio (w/w). Vortex to ensure thorough mixing.
10. Incubate the samples for digestion for 3 hours at 37°C (0 rpm).
11. Quench the digestion reaction by adding trifluoroacetic acid (TFA) to a 1% final concentration.

**3. Peptide Clean-up (Pierce™ Peptide Desalting Spin Columns)**

12. Column Preparation: Remove the bottom tip, place the column into a 2 mL microcentrifuge tube, and centrifuge at 5,000 × g for 1 minute to pack the resin. Discard the liquid.
13. Conditioning: Wash the resin twice with 300 µL of ACN, centrifuging at 5,000 × g for 1 minute after each wash.
14. Equilibration: Wash the column twice with 300 µL of 0.1% TFA in water. The column is now conditioned.
15. Sample Loading: Load 300 µL of the sample solution onto the column and centrifuge at 3,000 × g for 1 minute. Discard the flowthrough.
16. Washing: Wash the column three times with 300 µL of 0.1% TFA solution, centrifuging at 3,000 × g for 1 minute after each wash. Discard the eluate after each wash.
17. Elution: Place the column into a new 2 mL microcentrifuge tube.
18. Elute the peptides by applying 300 µL of the 50% ACN, 0.1% TFA solution and centrifuging at 3,000 × g for 1 minute.
19. Repeat the elution step once more with 300 µL of the elution solution, collecting the eluate into the same tube.

**4. Final Sample Preparation for LC-MS**

20. Evaporate the liquid contents of the collected eluate to dryness using vacuum centrifugation.
21. Re-suspend the dry peptide samples in an appropriate volume of 0.1% formic acid (FA) for LC-MS analysis.
22. The optimal peptide amount to be loaded onto the LC-MS system is 2 µg.

**B. Sample Preparation by Easy Extraction and Digestion (SPEED Method)**

**1. Protein Extraction and Lysis**

1. Resuspend the cell pellet in high-purity trifluoroacetic acid (TFA) at a 1:4 (v/v) sample-to-TFA ratio. Mix until homogenous and incubate at room temperature (RT) for 5 minutes.
2. Neutralize the lysate by adding 2 M Tris base stock solution at 8× the volume of the TFA used for lysis. Vortex thoroughly.
3. Perform a protein concentration assay, if necessary, and aliquot 50 µg of protein for subsequent steps.

**2. Reduction and Alkylation**

4. Bring the solution to a concentration of 10 mM TCEP (tris(2-carboxyethyl)phosphine) and 40 mM CAA (2-chloroacetamide).
5. Incubate the sample in a thermomixer at 95°C for 5 minutes.
6. Remove the sample and dilute 1:5 with Milli-Q water.

**3. Enzymatic Digestion**

7. Add the trypsin/lys-c protease mix based on a 1:20 enzyme-to-protein ratio (w/w). Vortex to ensure thorough mixing.
8. Incubate for protein digestion for 3 hours at 37°C (0 rpm in a thermomixer).
9. Stop the digestion by adding TFA to achieve a 2% final concentration.

**4. Peptide Clean-up (Pierce™ Peptide Desalting Spin Columns)**

10. Column Preparation: Remove the bottom tip, place the column into a 2 mL microcentrifuge tube, and centrifuge at 5,000 × g for 1 minute to pack the resin. Discard the liquid.
11. Conditioning: Wash the resin twice with 300 µL of ACN, centrifuging at 5,000 × g for 1 minute after each wash.
12. Equilibration: Wash the column twice with 300 µL of 0.1% TFA in water. The column is now conditioned.
13. Sample Loading: Load 300 µL of the sample solution onto the column and centrifuge at 3,000 × g for 1 minute. Discard the flowthrough.
14. Washing: Wash the column three times with 300 µL of 0.1% TFA solution, centrifuging at 3,000 × g for 1 minute after each wash. Discard the eluate after each wash.
15. Elution: Place the column into a new 2 mL microcentrifuge tube.
16. Elute the peptides by applying 300 µL of the 50% ACN, 0.1% TFA solution and centrifuging at 3,000 × g for 1 minute.
17. Repeat the elution step once more with 300 µL of the elution solution, collecting the eluate into the same tube.

**5. Final Preparation for LC-MS**

18. Evaporate the liquid contents of the collected eluate to dryness using vacuum centrifugation.
19. Re-suspend the dry peptide samples in an appropriate volume of 0.1% formic acid (FA) for LC-MS analysis.
20. The optimal peptide amount to be loaded onto the LC-MS system is 2 µg.
